# SH2-mediated steric occlusion of the C2 domain regulates autoinhibition of SHIP1 inositol 5-phosphatase

**DOI:** 10.1101/2025.09.02.673847

**Authors:** Emma E. Drew, Hunter G. Nyvall, Matthew A.H. Parson, Reed K. Talus, John E. Burke, Scott D. Hansen

## Abstract

The Src homology 2 (SH2) domain containing inositol polyphosphate 5-phosphatase 1 (SHIP1) is an immune cell specific enzyme that regulates phosphatidylinositol-(3,4,5)-trisphosphate signaling at the plasma membrane following receptor activation. SHIP1 plays an important role in processes such as directed cell migration, endocytosis, and cortical membrane oscillations. Alterations in SHIP1 expression have been shown to perturb myeloid cell chemotaxis and differentiation. In the brain, SHIP1 regulate microglial cell behaviors, which has been linked to Alzheimer’s disease. Understanding the structural and functional relationships of SHIP1 is critical for developing ways to modulate SHIP1 membrane localization and lipid phosphatase activity during immune cell signaling. Recently, we discovered that the N-terminal SH2 domain of SHIP1 suppresses lipid phosphatase activity. SHIP1 autoinhibition can be relieved through interactions with receptor-derived phosphotyrosine (pY) peptides presented on membranes or in solution. Using hydrogen-deuterium exchange mass spectrometry (HDX-MS) we identified intramolecular contacts between the N-terminal SH2 domain and CBL1 motif of the C2 domain that limit SHIP1 membrane localization and activity. Single molecule measurements of purified SHIP1 on supported lipid bilayers and in neutrophil-like cells support a model in which the SH2 domain blocks membrane binding of the central catalytic module. Mutations that disrupt autoinhibition enhance the membrane binding frequency and increase the catalytic efficiency of SHIP1. Although dimerization of SHIP1 enhances membrane localization and the apparent phosphatase activity, it is not required for SHIP1 autoinhibition. Overall, our results provide new insight concerning SHIP1’s structural organization, membrane binding dynamics, and the mechanism of autoinhibition.

## INTRODUCTION

Phosphatidylinositol phosphate (PIP) lipids play a critical role in localizing proteins to intracellular membranes in eukaryotic cells (1, 2). The concentration and spatial distribution of PIP lipids is regulated by numerous lipid kinases and phosphatases that compete to establish unique steady-state concentrations required for cellular homeostasis and dynamic membrane signaling responses (3). In the context of cell polarity and migration pathways, receptor activation enhances the plasma membrane localization and activity of phosphoinositide 3-kinase (PI3K), which drives the rapid phosphorylation of phosphatidylinositol-(4,5)-bisphosphate (PI(4,5)P_2_) to generate phosphatidylinositol-(3,4,5)-trisphosphate (PI(3,4,5)P_3_) (4–7). Functioning downstream of the PI3K signaling pathway, the hematopoietic specific Src homology 2 (SH2) domain containing inositol polyphosphate 5-phosphatase 1 (SHIP1) plays a crucial role in regulating the dephosphorylation of PI(3,4,5) P_3_ to produce PI(3,4)P_2_ (8–10). Genetic knockout (KO) or silencing of SHIP1 expression perturbs cell polarity and migration in neutrophils (11–14). SHIP1 knockout mice develop chronic myelogenous leukemia characterized by the hyperproliferation and infiltration of myeloid cells in lung tissue (15). In the nervous system, SHIP1 is highly expressed in microglial cells where it regulates the inflammasome (16, 17). Misregulation of SHIP1 and microglial cell function are a marker for Alzheimer’s disease (17, 18). Defining the mechanisms that regulate SHIP1 activation is critical for deciphering how this lipid 5-phosphatase controls myeloid cell signaling.

SHIP1 is a 145 kDa multidomain protein that is predicted to lack secondary structure across ∼30% of its primary amino acid sequence (**Figure 1A**). The intrinsically disordered C-terminal domain (872-1189 aa) contains multiple NPxY and proline-rich motifs that can interact with phosphotyrosine binding (PTB) and SH3 domain containing proteins, respectively (19). Positioned in the mostly unstructured peptide sequence connecting the N-terminal SH2 domain and the PH-R domain (**Figure 1A**), the RhoA binding domain (RBD) has been shown to regulate dimerization of the SHIP1 paralog, SHIP2 (20). Flanking either side of the central 5-phosphatase domain are pleckstrin homology (PH) related and C2 domains (**Figure 1A**), which display weak affinities for PI(3,4,5)P_3_ and PS lipids, respectively (21–24). X-ray crystallography data indicates that the C2 domain forms multiple contacts with the 5-phosphatase domain (21, 23). In the case of SHIP2, the C2 domain modulates the activity of the phosphatase domain, which has led researchers to propose a model of allosteric regulation (23). The residues that bridge the phosphatase and C2 domain interface, however, are not conserved in SHIP1 (21).

**Figure 1.**
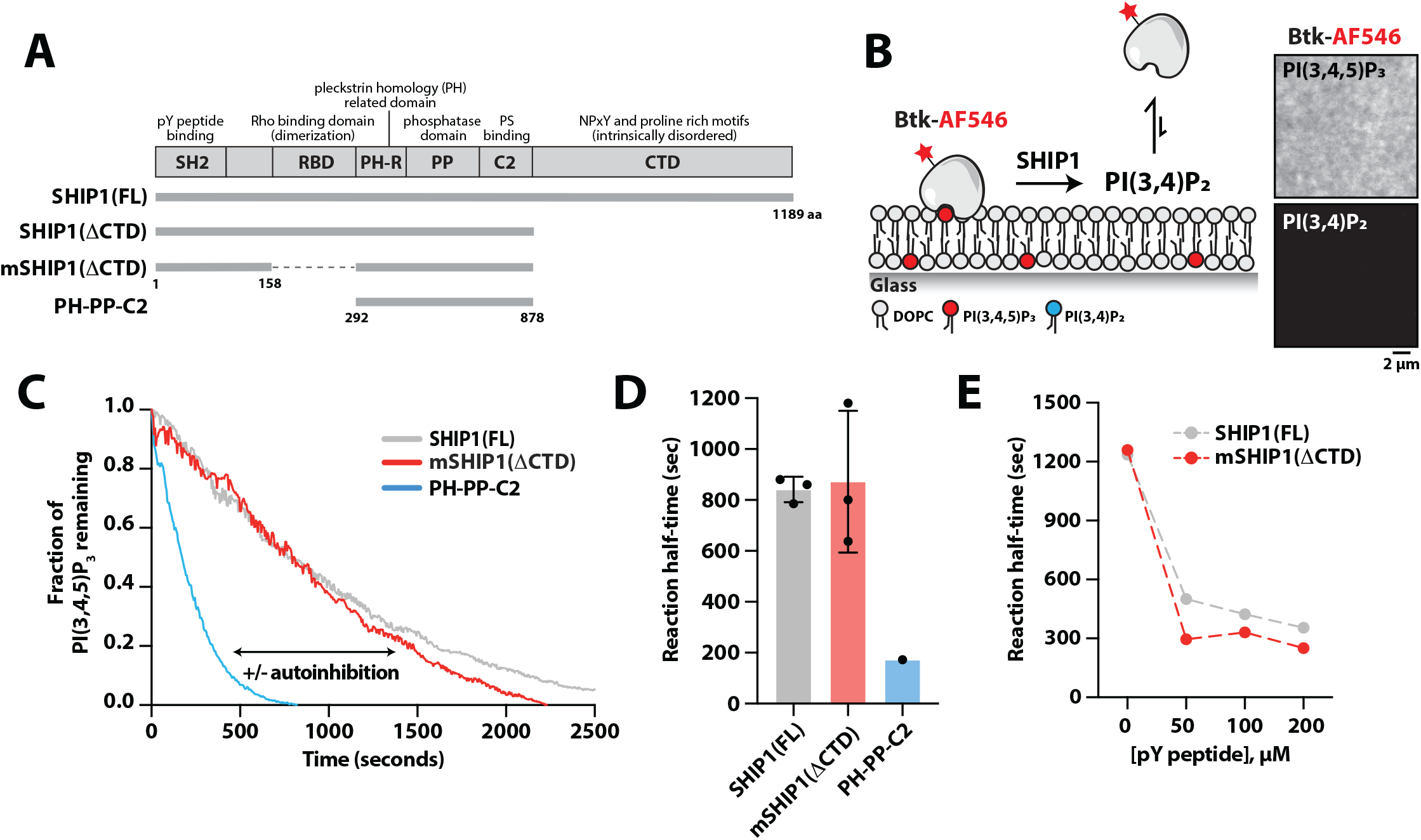
A minimal SHIP1 protein that exhibits SH2-dependent autoinhibition. **(A)** Domain organization of SHIP1 proteins biochemically characterized with indicated amino acid numbers. The dashed line represents residues in the Rho binding domain (RBD) that are absent in monomeric SHIP1(ΔCTD). **(B)** Diagram of the experimental design of in vitro SLB experiments to measure SHIP1 activity. Fluorescently labeled Btk1 binds specifically to PI(3,4,5)P_3_ and dissociates from the membrane when SHIP1 dephosphorylates PI(3,4,5)P_3_ to PI(3,4)P_2_. Inset images show representative TIRF-M images of the localization of 20 nM Btk-AF546 on 2% PI(3,4,5)P_3_ or 2% PI(3,4)P_2_. **(C)** Kinetic traces of SHIP1 phosphatase activity in the presence of 20 nM mNG-SHIP1 (FL), 20 nM mNG-mSHIP1(ΔCTD), or 20 nM mNG-SHIP1(PH-PP-C2). **(D)** Quantification of mean reaction half-time measured in the presence of 20 nM mNG-SHIP1(FL), mNG-mSHIP1(ΔCTD), or mNG-SHIP1(PH-PP-C2). N = 3 technical replicates per construct, p=0.868 comparing SHIP1(FL) and mSHIP1(ΔCTD). **(E)** Tyrosine phosphorylated peptides (pY) stimulate the activation of both mNG-mSHIP1(ΔCTD) and mNG-SHIP1(FL) in solution. For all reactions, dephosphorylation of PI(3,4,5)P_3_ was monitored in the presence of 20 nM Btk-SNAP-AF546. **(C-E)** Initial membrane composition: 2% PI(3,4,5)P_3_, 98% DOPC. Bars represent means. Errors equal SD. P-values calculated by student t-test.

Previously, we reported that full-length SHIP1 exhibits autoinhibition based on an undefined set of interactions that depend on the N-terminal SH2 domain (24). Although tyrosine phosphorylated peptides derived from an immunoreceptor tyrosine inhibitory motif (ITIM) can stimulate SHIP1 membrane binding and activity (24), the mechanism of autoinhibition has remained unclear. Here, we present a combination of single molecule TIRF microscopy (smTIRF-M) and hydrogen deuterium exchange mass spectrometry (HDX-MS) experiments that elucidates the mechanism of SHIP1 autoinhibition. Solution-based HDX-MS experiments performed in the absence and presence of tyrosine phosphorylated peptides revealed changes in solvent exposure in the N-terminal SH2 and C2 domains, consistent with this being the auto-inhibitory interface. Comparing the deuterium exchange rates of monomeric and dimeric SHIP1 revealed similar pY-mediated conformational changes that were limited to the SH2 and C2 domains. Based on the consensus of multiple HDX-MS experiments, we performed mutational analysis to isolate SHIP1 mutants that lack autoinhibition. Similar to the increase in activity observed in the presence of pY peptides, mutations in the CBL1 motif of the C2 domain enhance the membrane binding frequency and increase the catalytic efficiency of SHIP1. Although dimerization enhances SHIP1 membrane localization, we find that a minimal construct lacking the dimerization domain and the unstructured C-terminal domain, retains the characteristics of the autoinhibited enzyme. Together, our results support a model in which the SH2 domain limits SHIP1 activity by sterically occluding membrane binding of the C2 domain.

## RESULTS

### A minimal SHIP1 protein that exhibits SH2-mediated autoinhibition

Previous biochemical characterization established that SHIP1 autoinhibition is mediated by an undefined set of interactions between the N-terminal SH2 domain and some portion of the central globular domains consisting of the pleckstrin homology (PH) related domain, phosphatase domain, and C2 domain **(Figure 1A)** (24). Due to the large fraction of intrinsically disordered peptide sequence, purification of full-length SHIP1 is challenging and less amenable to structural biochemistry required to decipher the mechanism of autoinhibition. Given the complexity of the full-length protein, we aimed to create a minimal SHIP1 construct that exhibits autoinhibition and can be activated with tyrosine phosphorylated (pY) peptides in solution or on supported membranes. Compared to full-length SHIP1, our minimal SHIP1 construct lacks the intrinsically disordered C-terminal domain (CTD). The CTD contributes a minor role to autoinhibition compared to the N-terminal SH2 domain (24). In addition, the minimal SHIP1 construct contains a shorter linker between the SH2 domain and the central domains (**Figure 1A**). Reducing the linker length removes the RhoA binding domain (RBD), which is predicted to dimerize based on homology to SHIP2 (20). Supporting this model, AlphaFold predicts that the SHIP1(RBD) dimerizes with high confidence (**Figure S1A-S1B**). In addition, purified SHIP1(RBD) fused to an mNeonGreen (mNG) eluted from a size exclusion chromatography column as a single peak with an apparent molecular weight of a dimer (**Figure S1C-S1D**). This was observed across multiple protein concentrations, consistent with the isolated RBD forming a constitutive dimer that does not readily dissociate into monomeric species when diluted.

To characterize the phosphatase activity of monomeric SHIP1 (mSHIP1) lacking the C-terminal domain (ΔCTD), we used a previously established in vitro supported lipid bilayer (SLB) assay to visualize the dephosphorylation of PI(3,4,5)P_3_ by total internal reflection fluorescence microscopy (TIRF-M) (24). Serving as a mimetic to the cellular plasma membrane but containing a more simplified lipid composition, these SLBs are fluid and miscible at room temperature. To measure the kinetics of PI(3,4,5)P_3_ dephosphorylation in the presence of recombinantly purified SHIP1, we used a fluorescently labeled lipid binding domain derived from Bruton’s tyrosine kinase (Btk) as a biosensor (**Figure 1B**). Compared to the Grp1-derived PI(3,4,5)P_3_ sensor used in our previous study (24), here we used a monomeric Btk mutant protein that interacts with a single PI(3,4,5)P_3_ lipid. This biosensor exhibits more rapid equilibration kinetics and a faster dissociation rate constant compared to Grp1 (25, 26). Using this assay, we measured the lipid phosphatase activity of full-length SHIP1 and mSHIP1(ΔCTD) (**Figure 1C-1D**). Both proteins exhibited ∼5-fold reduction in phosphatase activity compared to the central catalytic module of SHIP1 alone, which we refer to as PH-PP-C2 (**Figure 1C-1D**). The difference in activity between mSHIP1(ΔCTD) and constitutively active SHIP1 (PH-PP-C2) construct set the dynamic range for characterizing SHIP1 mutants that modulate lipid phosphatase activity. The addition of a pY peptide derived from an immunoreceptor tyrosine inhibitory motif (ITIM) of FCγRIIB in solution relieved autoinhibition and stimulated the phosphatase activity to similar extents for both full-length SHIP1 and mSHIP1(ΔCTD) (**Figure 1E**).

### HDX-MS reveals pY peptide mediated conformational changes in autoinhibited SHIP1

The structure of full-length autoinhibited SHIP1 has not been solved by X-ray crystallography or cryo-electron microscopy. To determine the structural basis of SHIP1 autoinhibition, we performed hydrogen deuterium exchange mass spectrometry (HDX-MS) experiments to identify peptide sequences that regulate intramolecular interactions between the N-terminal SH2 domain and presumably the central catalytic core flanked by the PH and C2 domains. HDX-MS measures the exchange rate of amide hydrogens acting as a readout of protein secondary structure dynamics. HDX-MS is capable of identifying both direct and allosteric changes which occur in the presence of ligand (27). We performed solution experiments with mSHIP1(ΔCTD) in the absence and presence of pY-ITIM peptides (**Figure 2A**). The HDX-MS experiments identified SHIP1 peptides in the SH2 domain that exhibited both increased exchange (46-52aa) and decreased exchange (residues 29-39aa and 64-72aa), as well as peptides in the C2 domain that displayed an increase in exchange in the presence of the pY-ITIM peptide (**Figure 2A-2B** and **Figure S2A**). In particular, the disordered CBL1 motif in the C2 domain (740-748aa) displayed increased solvent exposure in the presence of pY-ITIM peptides.

**Figure 2.**
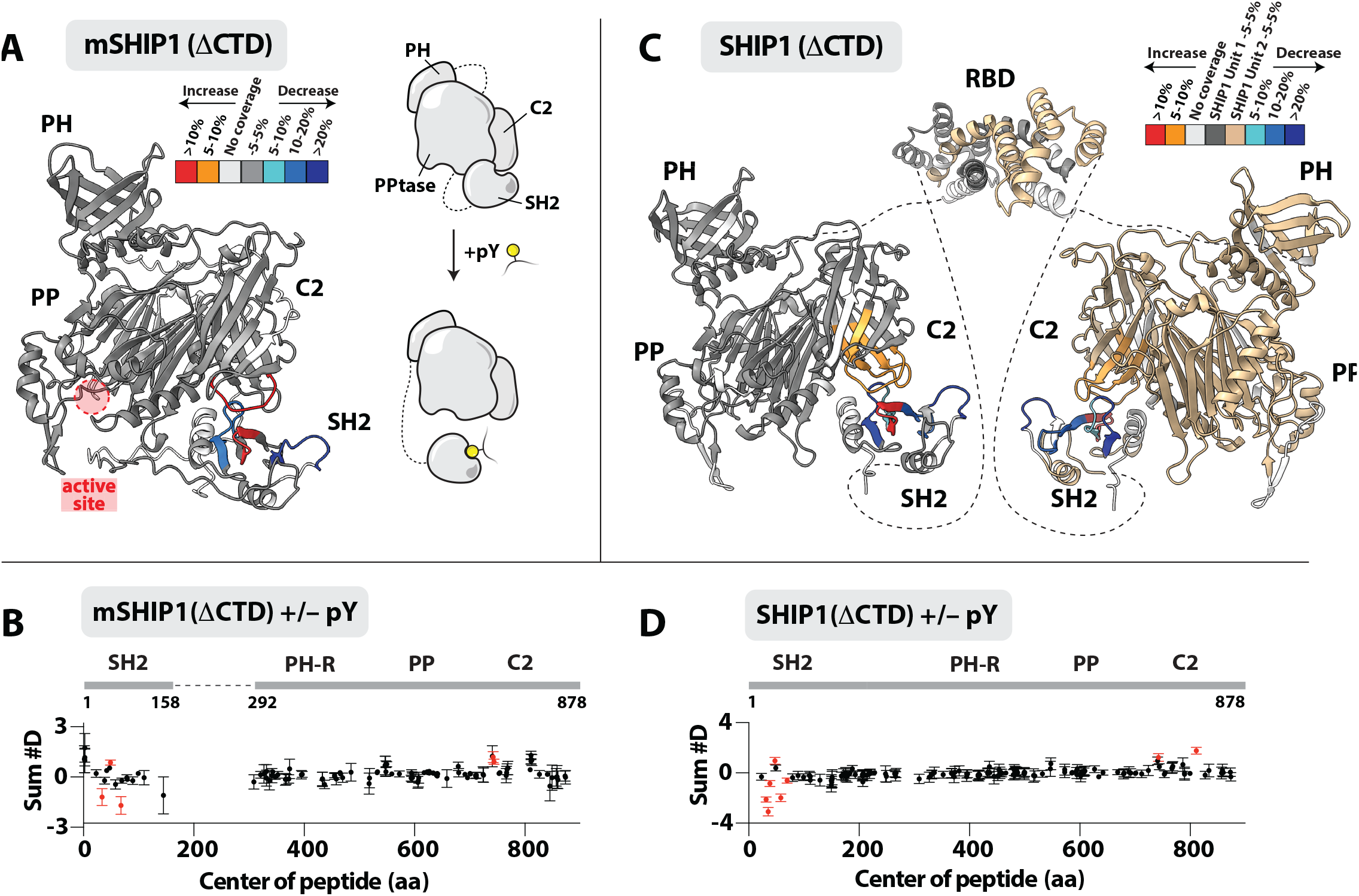
HDX-MS reveals intramolecular contacts between the SH2 and C2 domain of SHIP1. **(A)** Peptides in the SH2 and C2 domains of monomeric SHIP1(ΔCTD) (1.5 µM) showing significant changes in HDX (>0.4 Da and 5% difference, with a two-tailed t-test p < 0.01) when incubated with 40 µM pY peptide are mapped onto an AlphaFold2 model for monomeric mSHIP1(ΔCTD) (also refer to **Figure S3**). Differences in exchange are color coded according to the legend. Cartoon model summarizing pY-mediated conformational changes in mSHIP1(ΔCTD) is shown to the right. **(B)** Sum of the number of deuteron difference in deuterium incorporation for mSHIP1(ΔCTD) measured in the absence or presence of 40 µM pY peptide. **(C)** Autoinhibited SHIP1(ΔCTD) (1.5 µM) showing significant changes in HDX (>0.4 Da and 5% difference with a two-tailed t-test p < 0.01) when incubated with 40 µM pY peptide are mapped onto an AlphaFold3 model for dimeric SHIP1. Dimerization of the Rho binding domain (RBD) was inferred from AlphaFold3 structure prediction. Generation of dimer model from AlphaFold3 output is shown in **Figure S4**. Dashed lines represent linkers that are predicted to lack secondary structure. Differences in exchange are color coded according to the legend. **(D)** Sum of the number of deuteron difference in deuterium incorporation for SHIP1(ΔCTD) measured in the absence or presence of 40 µM pY peptide. **(B, D)** Values are based on the integrated difference in exchange over the entire time course. Each point represents a single peptide and error bars are shown as the sum of the standard deviation across all time points (n = 3 for each time point). Peptides with a statistically significant difference in HDX are highlighted in red, with the domain schematic annotated above. See **Figure S2** for a more complete set of peptides comparing apo mSHIP1(ΔCTD) and SHIP1(ΔCTD) deuterium incorporation in the absence and presence of pY peptide. Refer to ***Methods*** and **Tables S1-S2** for full description of statistical evaluation method.

In agreement with our HDX-MS data, AlphaFold (28, 29) consistently predicted an autoinhibited mSHIP1(ΔCTD) structure with potential intramolecular contacts between the SH2 and C2 domains (**Figure 2A** and **Figure S3A-B**). The predicted structure for the SHIP1 (PPtase-C2) was in good agreement with the SHIP1 and SHIP2 (PPtase-C2) structures previously solved by X-ray crystallography (21, 23) (**Figure S3C**).

To determine whether the SHIP1 dimerization motif modulates pY peptide mediated solvent exposure of the SH2 and C2 domains, we performed HDX-MS experiments using SHIP1(ΔCTD) (**Figure 2C**). This construct lacks the disordered C-terminal domain but contains the RBD dimerization domain (152-291 aa). Using AlphaFold3 (30), we mapped pY peptide dependent changes in hydrogen deuterium exchange to a model of SHIP1(ΔCTD) in a dimeric state (**Figure 2C-2D** and **Figure S4**). Overall, the HDX-MS experiments performed with SHIP1(ΔCTD) identified intramolecular contacts between the SH2 and the CBL1 motif in agreement with our results using the mSHIP1(ΔCTD) construct lacking the RBD (**Figure 2A**). In addition to observing solvent exposure of the CBL1 motif, the HDX-MS results also revealed a second loop exposure in the CBL3 motif of the C2 domain (**Figure 2C-2D**). Note that the AlphaFold3 structure prediction tries to maximize the surface area of inter-subunit contact in the SHIP1 dimer (**Figure S4A**), particularly in the C2 and SH2 domains. However, the predicted alignment error (PAE) for dimer interfaces outside of the RBD was predicted with very low confidence, with a high confidence PAE prediction for the RBD dimer (**Figure S4B**). In addition, there is no structural biochemistry to support the proposed AlphaFold3 intramolecular interactions beyond those mediated by the dimerization domain (**Figure S4B-S4D**). As a result, our structural model assumes that the RBD is the only point of contact between two SHIP1 subunits in a dimer (**Figure 2C** and **Figure S4D**).

### Mutations in the C2 domain disrupt SHIP1 autoinhibition

Based on our HDX-MS data, we hypothesized that the SH2 domain binds to the C2 domain, which limits SHIP1 interactions with membrane lipids such as phosphatidylserine (PS). Intramolecular interactions between the SH2 and C2 domains could also prevent the ability of the C2 domain to change conformation, which has been proposed to allosterically regulate 5-phosphatase activity of SHIP2 (23). To determine the functional significance of the SH2-C2 intramolecular interactions, we purified mSHIP1(ΔCTD) containing mutations in the conserved CBL1 motif (**Figure 3A**). The SHIP1 CBL1 motif contains multiple positively charged lysine residues that could mediate electrostatic interdomain interactions (**Figure 3A-3B**). We purified mSHIP1(ΔCTD) mutants with the lysine residues mutated to uncharged alanine residues (K741A/K743A/K747A; denoted 3A) or negatively charged aspartic acid residues (K741D/K743D/K747D; denoted 3D) (**Figure 3C**). In addition, we mutated all the residues in the disordered CBL1 motif to alanine residues (denoted 7A) (**Figure 3C-3D**). Using our SLB TIRF microscopy assay, we measured the phosphatase activity of the SHIP1 CBL1 mutants compared to autoinhibited mSHIP1(ΔCTD) and the constitutively active central domain (i.e. PH-PP-C2) (**Figure 3E**). Mutating three CBL1 motif residues to uncharged residues (i.e. 3K to 3A) did not alter the phosphatase activity of mSHIP1(ΔCTD) (**Figure S5A-S5C**). In contrast, the mSHIP1(ΔCTD) containing the 3D charge swap or 7A loop mutant displayed increased 5-phosphatase activity with kinetic traces indistinguishable from constitutively active SHIP1 (PH-PP-C2) (**Figure 3E-3F**).

**Figure 3.**
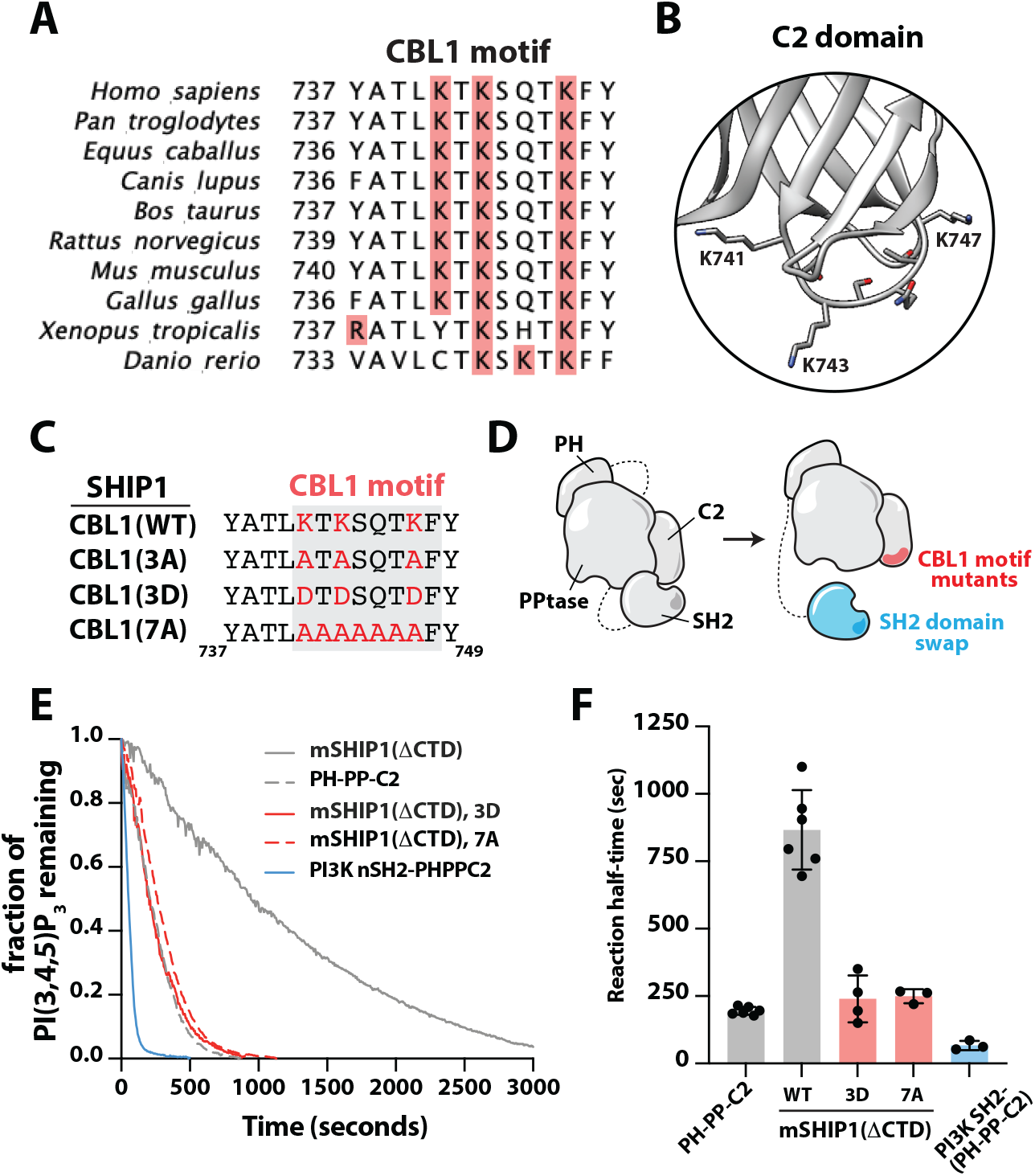
Mutations in the SH2 domain or CBL1 motif disrupt SHIP1 autoinhibition. **(A)** Sequence alignment of CBL1 motif across vertebrate species. Highlighted in red are positively charged residues within the CBL1 motif. **(B)** Zoom in structural model of human SHIP1 CBL1 motif with indicated lysine residues (PDB: 6IBD). **(C)** Sequence alignment summarizing human SHIP1 CBL1 motif mutant generated in this study. **(D)** Cartoons of SHIP1 autoinhibition mutants. **(E)** SHIP1 CBL1 motif mutants and SH2 domain swap display increased phosphatase activity. Kinetic traces of phosphatase activity measured in the presence of 20 nM mNG-mSHIP1(ΔCTD) (WT and mutants). **(F)** Quantification of reaction half-times of 20 nM mNG-SHIP1(PH-PP-C2), mNG-mSHIP1(ΔCTD), and mNG-mSHIP1(ΔCTD) mutants (N=3-6 technical replicate per construct). Statistical significance evaluated by student t-test comparing mSHIP1(ΔCTD) to the following mutants: 3D (*p<0.0001), 7A (*p<0.0001), PI3K(SH2)-SHIP1(PH-PP-C2) (*p<0.0001). Bars represent mean. Errors equal SD. P-values calculated by student t-test. **(E-F)** Initial membrane composition: 2% PI(3,4,5)P_3_, 98% DOPC. For all SHIP1 phosphatase activity assays, dephosphorylation of PI(3,4,5)P_3_ was monitored in the presence of 20 nM Btk-SNAP-AF546.

The AlphaFold prediction of autoinhibited mSHIP1(ΔCTD) did not reveal an obvious patch of negatively charged residues on the SH2 domain that could facilitate interactions with the CBL1 motif of the C2 domain (**Figure 2C** and **Figure S4**). Interpretating the HDX-MS data is also challenging because pY peptide binding can mask SH2 domain solvent exposure when SHIP1 autoinhibition is relieved. SH2 domain residues shielded by the pY peptide could also be responsible for both SHIP1 autoinhibition and pY peptide binding. To determine the role of the SH2 domain in regulating autoinhibition, we instead made a chimeric protein containing the nSH2 domain derived from human PI3Kα fused to SHIP1 (PH-PP-C2) in the same molecular configuration of mSHIP1(ΔCTD) (**Figure 3D**). Compared to our minimal construct, mSHIP1(ΔCTD), the PI3Kα(nSH2)-PH-PP-C2 chimera displayed enhanced 5-phosphatase activity (**Figure 3E-3F**). Interestingly, the catalytic activity of the chimera was slightly greater than the 3D and 7A mutants that disrupt SHIP1 autoinhibition. Based on our single molecule TIRF-M measurements, the increase in activity can be explained by a 2.6-fold increase in membrane binding frequency for PI3Kα(nSH2)-PH-PP-C2 compared to the SHIP1 (PH-PP-C2) construct (**Figure S6A-S6B**).

### CBL1 motif mutants maintain sensitivity to phosphatidylserine lipids

Previous work established that phosphatidylserine (PS) lipids enhance SHIP1/2 phosphatase activity independent of autoinhibition (23, 24). Since C2 domains are known for binding PS lipids (31), we wanted to determine the functional significance of the CBL1 motif in controlling PS-mediated enhancement of SHIP1 phosphatase activity. Using the SLB TIRF assay to monitor phosphatase activity, we created bilayers with a physiologically relevant concentrations of 20% PS lipids to measure the activity of the CBL1 mutants (32). The activity of mSHIP1(ΔCTD), mSHIP1(ΔCTD, 3D), and SHIP1(PH-PP-C2) followed the same trend as activity measurements performed on bilayers in the absence of PS lipids **(Figure 4A-4B)**. However, these experiments used a 10-fold lower concentration of SHIP1 protein to produce similar reaction half-times measured in the absence of PS lipids. In agreement with previous research, the enhanced activity was independent of autoinhibition since constructs lacking the SH2 domain also exhibit increased catalytic efficiency.

**Figure 4.**
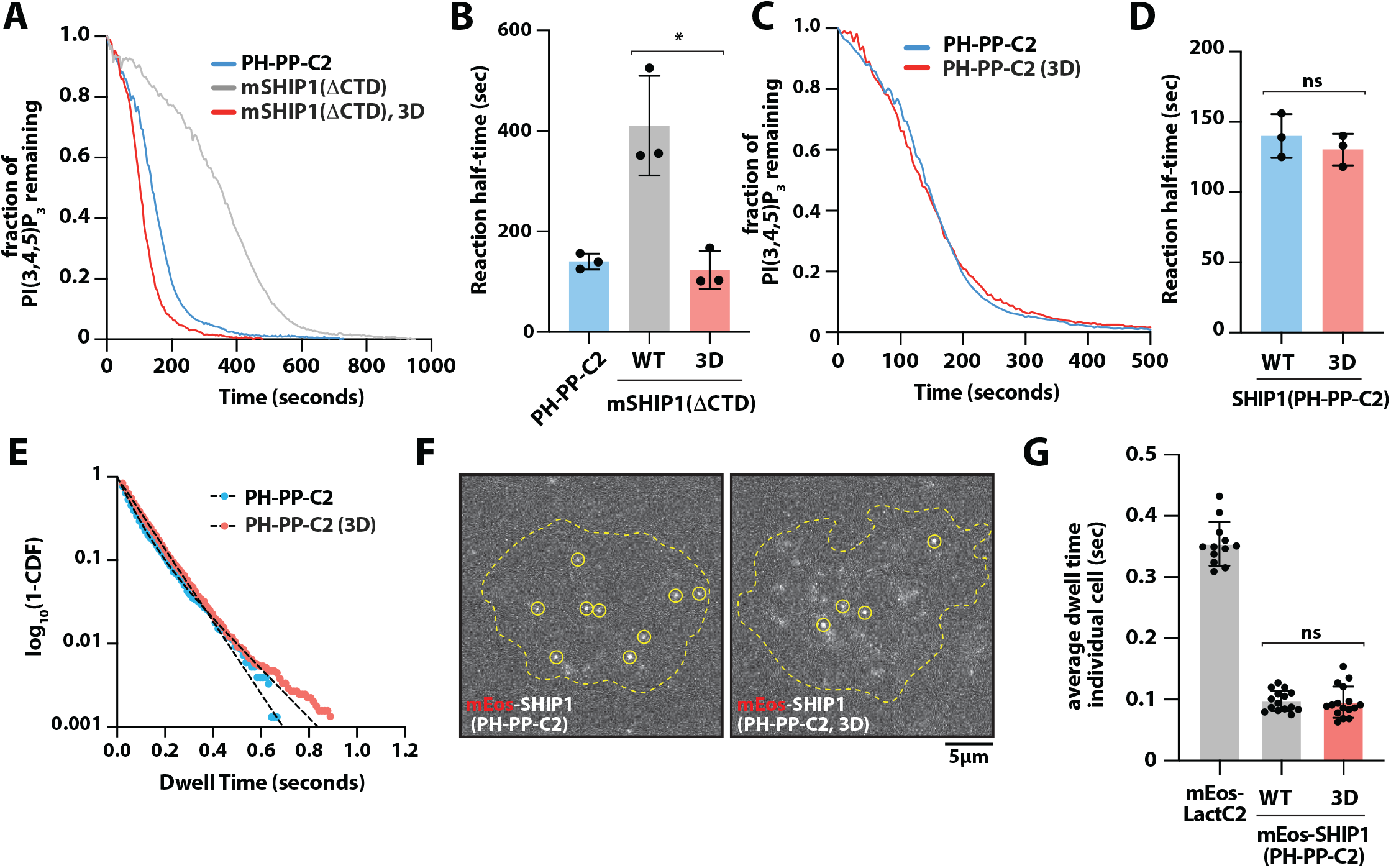
Phosphatidylserine lipids similarly enhance phosphatase activity of SHIP1 and CBL1 motif mutants. **(A)** Lipid phosphatase activity measured in the presence of 2 nM mNG-SHIP1 (PH-PP-C2) and mNG-mSHIP1(ΔCTD) (WT and mutants). **(B)** Quantification of reaction half-times of PH-PP-C2, mSHIP1(ΔCTD) and mSHIP1(ΔCTD) (K741D/K743D/K747D; denoted 3D). Bars equal mean reaction half-times. (N = 3 technical replicates per construct). p=0.009 comparing mSHIP1(ΔCTD) and mSHIP1(ΔCTD, 3D). **(C)** Phosphatase activity measurements of 2 nM mNG-SHIP1 (PH-PP-C2) WT and 3D mutant. **(D)** Quantification of reaction half-time of data in **(C)**. (N = 3 technical replicates per construct, p = 0.432, ns). **(E)** Single molecule dwell time distributions for all mNG-SHIP1 membrane binding events. Dwell times were calculated by fitting log_10_(1-cumulative distribution frequency (CDF)) to a single exponential decay curve (black dashed lines). **(F)** Representative TIRF-M images showing single molecule plasma membrane localization of indicated mEos-tagged SHIP1 construct in PLB-985 cells. **(G)** Average single molecule membrane dwell times for mEos-SHIP1(PH-PP-C2) (N=15 cells), mEos-SHIP1(PH-PP-C2, 3D) (N=16 cells), and mEos-LactC2 (N=12 cells). Each data point is the average dwell measured for an individual cell. p=0.806 comparing mEos-SHIP1(PH-PP-C2) and mEos-SHIP1(PH-PP-C2, 3D). **(A-E)** Membrane composition: 2% PI(3,4,5)P_3_, 20% DOPS, 78% DOPC. For all kinetic assays, dephosphorylation of PI(3,4,5)P_3_ was monitored in the presence of 20 nM Btk-SNAP-AF546. Errors equal SD. P-values calculated by student t-test.

Independent of intramolecular interactions between the SH2 and C2 domains, mutations in the CBL1 motif could stimulate 5-phosphatase activity through an allosteric mechanism based on altered interdomain communication between the central phosphatase and C2 domains. This would be surprising given that the CBL1 motif is positioned distal to residues that form the interface between the phosphatase and C2 domains to regulate allostery (21–23). To test whether CBL1 motif mutations modulate phosphatase activity independent of the propose autoinhibition mechanism, we introduced the CBL1 motif mutations (i.e. 3K to 3D) into the constitutively active SHIP1 construct (i.e. PH-PP-C2). When we measured the activity of SHIP1(PH-PP-C2) and SHIP1(PH-PP-C2,3D) in the presence of 2% PI(3,4,5)P_3_ and 20% PS lipids, we observed nearly identical kinetic traces and reaction half-times (**Figure 4C-4D)**. Consistent with this data, single molecule dwell time analysis comparing mNG-SHIP1 (PH-PP-C2), wild-type and 3D mutant, yielded dwell times that were indistinguishable (**Figure 4E** and **Table 1**). Lateral diffusion of membrane bound mNG-SHIP1 (PH-PP-C2), wild-type and 3D mutant, were also nearly identical (**Figure S7** and **Table 1**). This indicates that the charge swap mutations in the CBL1 motif do not affect the frictional drag coefficient or membrane contact surface area of the mNG-SHIP1(PH-PP-C2) protein. Overall, mutations in the CBL1 motif do not impact PS lipid binding of SHIP1 in vitro. When we measured SHIP1 activity on SLBs lacking PS lipids, however, we observed a 22% increase in the activity of the CBL1 mutant compared to the wild-type phosphatase (**Figure S5D-S5E**). This can be rationalized by the lack of 20% PS lipids in the SLBs, which reduces charge repulsion and causes a small amount of non-specific binding of the 3D CBL1 mutant. Importantly, the magnitude of this difference cannot explain the ∼3.5-fold enhancement in activity observed when the 3D or 7A CBL1 mutants when introduced into the autoinhibited mSHIP1(ΔCTD) construct (**Figure 3F**).

**Table 1.** Single molecule analysis of mNG-SHIP1 on supported lipid bilayers. *SD* = standard deviation from the indicated number of technical replicates. *N* = # of SLBs for calculating the mean dwell times (i.e. technical replicates). *n* = total number of molecules tracked across the indicated number of technical replicates (*N*). Alpha (α) = fraction of molecules with characteristic dwell time (τ_1_) or diffusion coefficient (*D1*). The fraction of long dwelling species (τ_1_) or fast diffusing molecules (*D1*) equals 1– α. *Steps* = total number of displacements measured for all single molecule trajectories from all technical replicates (*N*). Membrane composition for mNG-SHIP1(ΔCTD & FL): 2% PI(3,4,5)P_3_, 2% MCC-PE, 96% DOPC with pY conjugated peptide. Membrane composition for mNG-SHIP1(PH-PP-C2, WT and 3D): 2% PI(3,4,5)P_3_, 20% DOPS, 78% DOPC. Photobleaching kinetics of mNG-SHIP1(PH-PP-C2) immobilized on glass yielded a time constant (τ _bleach_) of 2.7 seconds.

| Protein visualized | $\tau_1 \pm SD$ (sec) | $\tau_2 \pm SD$ (sec) | $\alpha \pm SD$ | *N* | *n* |
| --- | --- | --- | --- | --- | --- |
| mNG-mSHIP1(ΔCTD) | 0.409 ± 0.009 | - | - | 3 | 12813 |
| mNG-SHIP1 (ΔCTD) | 0.359 ± 0.007 | 1.83 ± 0.03 | 0.74 ± 0.002 | 2 | 8451 |
| mNG-SHIP1 (ΔCTD), dimers | 1.79 ± 0.001 | - | - | 2 | 939 |
| mNG-SHIP1 (FL) | 0.333 ± 0.03 | 1.81 ± 0.08 | 0.49 ± 0.02 | 3 | 7888 |
| mNG-SHIP1 (FL), dimers | 2.11 ± 0.14 | - | - | 3 | 1680 |
| mNG-SHIP1 (PH-PP-C2) | 0.083 ± 0.011 | - | - | 4 | 13495 |
| mNG-SHIP1 (PH-PP-C2), 3D | 0.078 ± 0.011 | - | - | 4 | 18784 |
| Protein visualized | *D1* ± *SD* ($\mu\text{m}^2/\text{sec}$) | *D2* ± *SD* ($\mu\text{m}^2/\text{sec}$) | $\alpha \pm SD$ | *N* | *steps* |
| mNG-SHIP1 (PH-PP-C2) | 0.196 ± 0.015 | 1.61 ± 0.243 | 0.79 ± 0.03 | 4 | 110095 |
| mNG-SHIP1 (PH-PP-C2), 3D | 0.208 ± 0.025 | 1.98 ± 0.431 | 0.76 ± 0.02 | 4 | 139805 |
| mNG-mSHIP1(ΔCTD) | 0.065 ± 0.005 | 0.429 ± 0.004 | 0.31 ± 0.007 | 3 | 117808 |
| mNG-SHIP1 (ΔCTD) | 0.035 ± 0.003 | 0.226 ± 0.017 | 0.42 ± 0.009 | 2 | 125409 |
| mNG-SHIP1 (FL) | 0.028 ± 0.001 | 0.164 ± 0.003 | 0.34 ± 0.017 | 3 | 168117 |

To measure SHIP1 membrane binding properties on a more complex membrane, we expressed SHIP1 (PH-PP-C2) and SHIP1 (PH-PP-C2, 3D) mutant as mEos3.2 fusion proteins in PLB-985 neutrophil-like cells as previously described (24). Stable lentiviral-based expression resulted in a low-level of expression with uniform localization across the plasma membrane. To resolve single membrane bound molecules, a fraction of mEos-SHIP1 proteins were photoconverted from the green to red fluorescent state through transient exposure to 405 nm light **(Figure 4F)**. This allowed us to visualize single mEos-SHIP1 molecule and measure their membrane binding properties in vivo. Comparing mEos-SHIP1 (PH-PP-C2) and mEos-SHIP1 (PH-PP-C2, 3D) across numerous cells revealed no significant difference in the average membrane dwell time (**Figure 4G**). Consistent with our dwell time analysis of mNG-SHIP1(PH-PP-C2) on SLBs, dwell times measured on the plasma membrane of immune cells were nearly identical to our in vitro measurements (**Table 1**). To determine if our ability to measure differences between mEos-SHIP1 constructs was masked by a photobleaching artifact, we quantified the dwell time of plasma membrane localized mEos-LactC2. This lipid binding domain interacts strongly with phosphatidylserine and exhibited an average membrane dwell time ∼3.5 times greater than our mEos-SHIP1 constructs (**Figure 4G**).

### Autoinhibition reduces the membrane binding frequency of SHIP1

Based on our HDX-MS results and SHIP1 structure predictions, we hypothesize that the SH2-C2 interaction would partially occlude membrane binding in the presence of PS and PI(3,4,5)P_3_ lipids (**Figure 5A)**. To test for this mechanism, we measured the membrane binding frequency of various mNG tagged SHIP1 constructs on supported membranes using single molecule TIRF microscopy (**Figure 5B-5C**). Consistent with the SH2 blocking the C2 domain mediated interaction with PS lipids, the cumulative membrane binding frequency of autoinhibited mNG-mSHIP1(ΔCTD) was 4.7-fold lower compared to mNG-SHIP1(PH-PP-C2) measured at the same protein concentration (**Figure 5D** and **Figure S6**). By contrast, mNG-mSHIP1(ΔCTD) containing either the 3D or 7A mutations in the CBL1 motif displayed an increased number of membrane binding events compared to the wild-type enzyme **(Figure 5D)**. By measuring the membrane binding frequency over a range of protein concentrations, we calculated the association rate constant (*k*_*ON*_) for each mNG-SHIP1 construct (**Figure S6**). The mNG-mSHIP1(ΔCTD) construct has an 8.3-fold lower *k*_*ON*_ compared to mNG-SHIP1 (PH-PP-C2) (**Figure 5E**). When measuring the *k*_*ON*_ for mNG-mSHIP1(ΔCTD) 3D and 7A mutants we observed an intermediate association rate constants that were 3.3- and 3.4-fold higher compared to wild-type mNG-mSHIP1(ΔCTD). This indicates that the CBL1 motif mutations reduce the ability of the SH2 domain to sterically hinder the C2 domain from membrane binding. Since mNG-SHIP1 (PH-PP-C2) and mNG-mSHIP1(ΔCTD) differ in molecular weight and overall structure, they are predicted to have slightly different membrane binding frequency based on diffusion through solution. Importantly, the membrane binding frequency of the mNG-mSHIP1(ΔCTD) 3D and 7A mutants were nearly identical, indicating that the net charge of the CBL1 motif does not strongly influence the membrane binding frequency.

**Figure 5.**
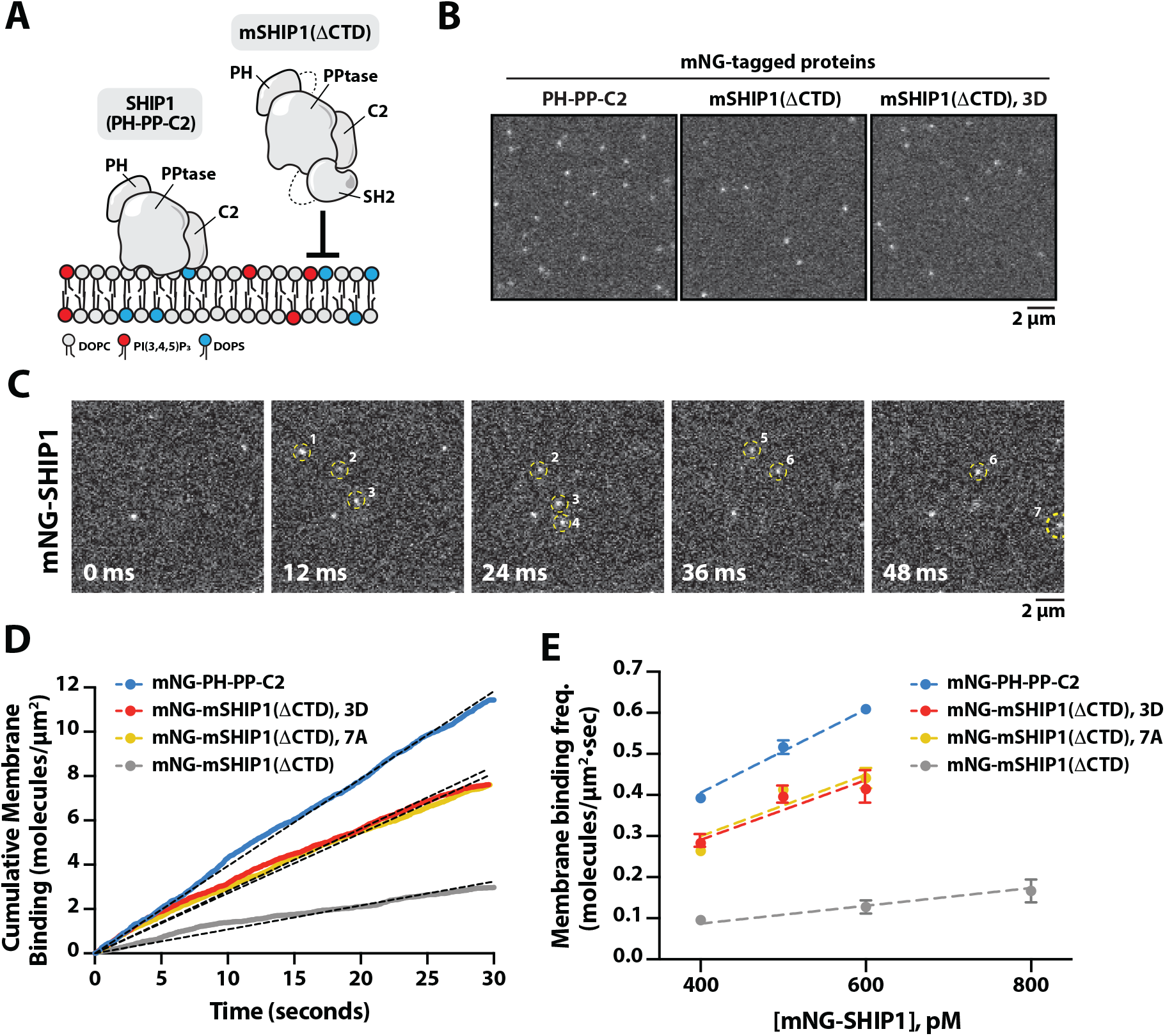
SH2-mediated steric occlusion of the CBL1 motif reduces the membrane binding frequency of SHIP1. **(A)** Cartoon illustrating predicted difference in membrane binding for SHIP1 (PH-PP-C2) and mSHIP1(ΔCTD) based SH2 mediated steric occlusion of the C2 domain. **(B)** Representative TIRF-M images of membrane localization of 400 pM mNG-SHIP1(PH-PP-C2), 400 pM mNG-mSHIP1(ΔCTD), and 400 pM mNG-mSHIP1(ΔCTD, K741D/K743D/K747D; denoted 3D). **(C)** Representative time sequence of images showing the frame-by-frame localization of mNG-SHIP1. New membrane binding events are scored and cumulated to calculate the binding frequency. **(D)** Cumulative binding events measured in the presence of 400 pM mNG-SHIP1 (PH-PP-C2), mNG-mSHIP1(ΔCTD), mNG-mSHIP1(ΔCTD, 3D), or mNG-mSHIP1(ΔCTD, 7A). **(E)** Membrane binding frequency (*k*_*ON*_) calculated from measuring cumulative binding events across the following protein concentrations: 400-600 pM mNG-SHIP1 (PH-PP-C2), 400-600 pM mNG-mSHIP1(ΔCTD, 3D), 400-600 pM mNG-mSHIP1(ΔCTD, 7A), and 400-800 pM mNG-mSHIP1(ΔCTD). Linear regression yielded the following *k*_*ON*_ values: 1.0 nM^-1^•µm^-2^•sec^-1^ mNG-SHIP1 (PH-PP-C2), 0.73 nM^-1^•µm^-2^•sec^-1^ mNG-mSHIP1(ΔCTD, 3D), 0.75 nM^-1^•µm^-2^•sec^-1^ mNG-mSHIP1(ΔCTD, 7A), and 0.22 nM^-1^•µm^-2^•sec^-1^ mNG-mSHIP1(ΔCTD). **(B-E)** Membrane composition: 2% PI(3,4,5)P_3_, 20% DOPS, 78% DOPC.

### Dimerization modulates SHIP1 membrane binding and apparent phosphatase activity

Since mSHIP1(ΔCTD) lacks the dimerization domain, we hypothesized that differences in oligomerization could modulate SHIP1 membrane localization and phosphatase activity. To mimic the physiological context of SHIP1 membrane signaling, we measured SHIP1 activity and localization on SLBs containing membrane conjugated pY-ITIM peptides (**Figure 6A**). For these experiments, we simultaneously monitored the dephosphorylation of PI(3,4,5)P_3_ and membrane density of various mNG-tagged SHIP1 proteins. The activity of mSHIP1(ΔCTD) was significantly reduced compared to full-length SHIP1 and the SHIP1(ΔCTD) mutant (**Figure 6B-6C**). Quantification of mNG fluorescent revealed that the difference in phosphatase activity could be explained by the enhanced membrane localization of mNG-SHIP1(FL) and mNG-SHIP1(ΔCTD), compared to mNG-mSHIP1(ΔCTD) (**Figure 6D**). Considering the difference in membrane localization between the various SHIP1 constructs, the phosphatase activity per SHIP1 molecule is nearly identical, while the apparent activities are significantly different at the same solution concentration.

**Figure 6.**
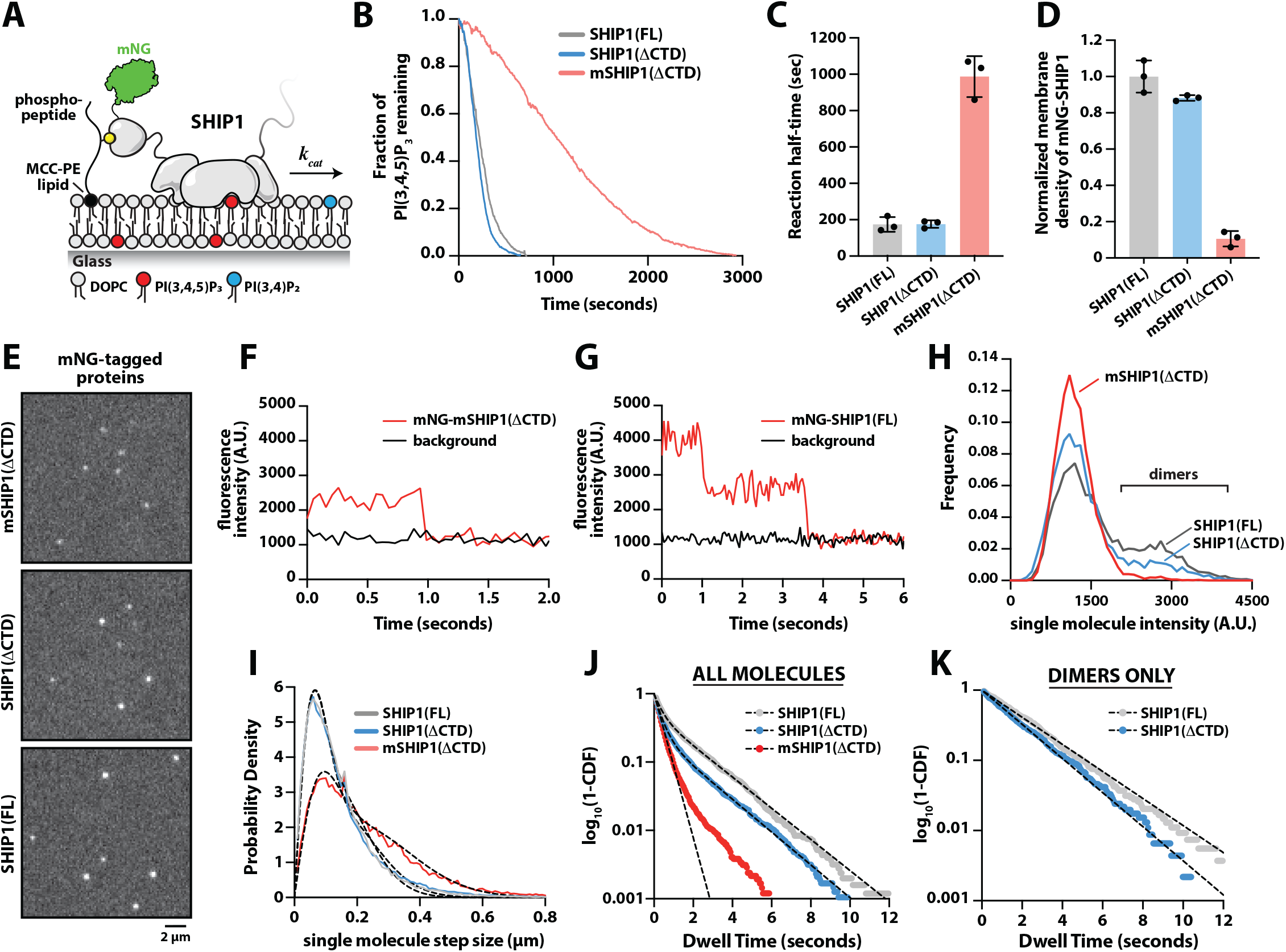
Dimerization enhances SHIP1 membrane recruitment and apparent phosphatase activity. **(A)** Cartoon illustrating experimental design for visualizing membrane recruitment of mNG-SHIP1 to pY peptides. **(B)** Representative kinetic traces of SHIP1 phosphatase activity in the presence of 100 pM mNG-SHIP1(FL), mNG-SHIP1(ΔCTD), or mNG-mSHIP1(ΔCTD). All reactions contained 20 nM Btk-SNAP-AF546 to measure dephosphorylation of PI(3,4,5)P_3._ **(C)** Quantification of reaction half-times for mNG-SHIP1(FL), mNG-SHIP1(ΔCTD), and mNG-mSHIP1(ΔCTD) in the presence of coupled pY-ITIM peptides (N=3 technical replicates per construct). **(D)** Normalized mNG-SHIP1 membrane surface density from reactions in (B). Fluorescence intensity of different mNG-SHIP1 constructs was normalized relative to the average membrane intensity of mNG-SHIP1(FL). **(E)** Representative TIRF-M imaging showing the single molecule localization of mNG-SHIP1 proteins (20-50 pM solution concentration). **(F-G)** Single molecule photobleaching dynamics of **(F)** mNG-mSHIP1(ΔCTD) and **(G)** mNG-SHIP1(FL) compared to background fluorescence (black line). **(H)** Molecular brightness distributions measured in the presence of 50 pM mNG-SHIP1(FL), 50 pM mNG-SHIP1(ΔCTD), or 20 pM mNG-mSHIP1(ΔCTD). **(I)** Step size distribution plots generated from single molecule tracking in the presence of either 50 pM mNG-SHIP1(FL), 50 pM mNG-SHIP1(ΔCTD), or 20 pM mNG-mSHIP1(ΔCTD). **(J)** Single molecule dwell time distributions for all mNG-SHIP1 membrane binding events. Dwell times were calculated by fitting log_10_(1-cumulative distribution frequency (CDF)) to either a single or double exponential decay curve (black dashed lines). **(K)** Single molecule dwell time distributions for all mNG-SHIP1 membrane binding events for only the legitimate dimers observed in the presence of either mNG-SHIP1(FL) or mNG-SHIP1(ΔCTD). **(B-K)** Initial membrane composition: 2% PI(3,4,5)P_3_, 2% MCC-PE (+pY-ITIM peptide), 96% DOPC. Refer to **Table 1** for statistics.

To quantify the abundance of SHIP1 dimers on membranes, we performed single molecule TIRF microscopy experiments to visualize mNG labeled SHIP1 (**Figure 6E**). We previously used this approach to quantify differences in the oligomerization state of type I phosphatidylinositol-4-phosphate 5-kinases (PIP5K) and a type II phosphatidylinositol-5-phosphate 4-kinase (PIP4K) bound to PI(4,5)P_2_ lipids (33). Using SLBs with conjugated pY peptides (**Figure 6A**), we performed single-molecule photobleaching experiments to determine the oligomerization state of membrane-bound mNG tagged SHIP1 proteins. In the case of mNG-mSHIP1(ΔCTD), membrane-bound particles photobleached in a single step (**Figure 6F**). By contrast, a subpopulation of mNG-SHIP1(FL) and mNG-SHIP1(ΔCTD) particles displayed two-step photobleaching events (**Figure 6G**). We then compared the single molecule brightness distributions for mNG-SHIP1 proteins bound to pY peptides. This analysis revealed the following fraction of legitimate dimers with two definitive mNG proteins visible: 1.7 ± 0.1% mNG-mSHIP1 (ΔCTD), 13 ± 1.4% mNG-SHIP1(ΔCTD), and 24 ± 1.8% mNG-SHIP1(FL) (**Figure 6H**). Considering that the degree of mNG chromophore maturation for recombinant proteins is approximately 75%, the maximum number of SHIP1 dimers we expect to observe is about 50% of the membrane bound particles. Since the fraction of observed SHIP1 dimers was lower than predicted for a constitutively dimeric protein, mNG-SHIP1 (FL) and mNG-SHIP1 (ΔCTD) appear to exist in a monomer-dimer equilibrium under our experimental conditions. If we assume that the membrane binding frequency of monomeric and dimeric mNG-SHIP1 are equivalent, we estimate that 48% of full-length mNG-SHIP1 and 26% of mNG-SHIP1 (ΔCTD) exists as dimers under the experimental conditions used for single molecule imaging (i.e. pM concentrations). Note that the fraction could be greater in the presence of a higher solution concentration of mNG-SHIP1.

Oligomerization of peripheral membrane binding proteins typically increases the frictional drag coefficient, which reduces membrane diffusivity (33– 35). Comparing the single molecule displacement across all trajectories revealed significantly slower membrane diffusivity for mNG-SHIP1(FL) and mNG-SHIP1(ΔCTD) compared to mNG-mSHIP1(ΔCTD) (**Figure 6I** and **Table 1**). Consistent with the RBD mediating dimerization, we also observed significantly longer membrane dwell times for mNG-SHIP1(FL) and mNG-SHIP1(ΔCTD) compared to mNG-mSHIP1(ΔCTD) (**Figure 6J** and **Table 1**). The dwell time distribution for mNG-mSHIP1(ΔCTD) could be fit using a single exponential (τ_1_ = 0.41 sec), while the SHIP1 constructs containing the RBD were best fit with a double exponential yielding two characteristic dwell times (τ_1_ and τ_2_; **Figure 6J** and **Table 1**). Interestingly, the fraction of long dwelling (τ_2_) mNG-SHIP1 (FL) and mNG-SHIP1 (ΔCTD) molecules was greater than the fraction of dimers calculated based on molecular brightness (**Table 1**; 1–α). As described above, this is consistent with our predicted number of dimers after considering the incomplete maturation of the mNG. Quantification of the dwell time of only legitimate mNG-SHIP1 (FL) and mNG-SHIP1(ΔCTD) dimers (i.e. two mNG visible) yielded a single time constant (τ_1_) similar to the long dwelling species (i.e. τ_2_) observed in the entire distribution of membrane binding events (**Figure 6K** and **Table 1**). Together, this suggests that the fraction of long dwelling mNG-SHIP1(FL and ΔCTD) molecules is a good proxy for the total number of dimer membrane binding events at equilibrium, which is 25-50%.

## DISCUSSION

To decipher the mechanism of SHIP1 autoinhibition, we created a minimal SHIP1 construct that displays the primary characteristics of full-length autoinhibited SHIP1. This is akin to the approach used to determine the molecular basis of N-WASP autoinhibition, coincidence detection, and activation in the context of actin filament nucleation (36, 37). Hydrogen deuterium exchange mass spectrometry (HDX-MS) experiments performed with mSHIP1(ΔCTD) revealed that the SH2 domain shields the CBL1/3 motifs in the C2 domain, suggesting a mechanism for autoinhibition. Importantly, similar interdomain interactions were detected in the absence and presence of the dimerization motif (i.e. Rho binding domain), which is positioned between the SH2 domain and central catalytic core (i.e. PH-PP-C2). Based on single molecule TIRF microscopy measurements of the membrane binding frequency, autoinhibition reduces the ability of SHIP1 to dock on membranes containing anionic lipids. Mutations in the CBL1 motif, including charge swaps (K741D/K743D/K747D) were sufficient to relieve autoinhibition and increase the phosphatase activity of mSHIP1(ΔCTD). Complete charge neutralization of the CBL1 motif (i.e. 7A mutation) similarly increased the activity and membrane binding of mSHIP1(ΔCTD), indicating the net charge of the CBL1 motif is not critical for membrane association. Our single molecule TIRF-M measurements indicate that mutations in the CBL1 motif increase the membrane binding frequency of SHIP1 by reducing steric hindrance by the SH2 domain. Although the SHIP1 CBL1 motif is slightly divergent from SHIP2, there is a conservation of basic residues which suggests a similar autoinhibitory mechanism exists in SHIP2.

Mutations in the CBL1 motif had no effect on the PS lipid enhanced activity measured in the context of the central catalytic core referred to as SHIP1(PH-PP-C2). In addition, CBL1 motif mutants did not perturb membrane dwell time and diffusivity of mNG-SHIP1(PH-PP-C2) measured on supported lipid bilayers and the plasma membrane of neutrophil-like cells. This suggests that the CBL1 motif modulates SHIP1’s activity through interactions with the SH2 domain and does not strongly influence PS binding. SHIP1/2 contains a P-variant (or Type-II) C2 domain, which is based on the orientation of Δ-sheets relative to the N- and C-terminal connections (38, 39). This naming convention, however, does not inform the function or specificity of the C2 domain. Unlike other lipid binding domains (e.g. pleckstrin homology domains), C2 domains generally lack a conserved binding pocket or basic residues that confer lipid specificity. C2 domains also have low primary amino acid sequence conservation (40). Although many type-II C2 domains bind calcium, SHIP1 and SHIP2 lack conserved acidic residues in their CBL motifs to coordinate calcium, which mediates PS lipid interactions. Consistent with these sequence characteristics, SHIP2 lipid binding and phosphatase activity is reportedly insensitive to calcium (23). Single molecule membrane binding measurements similarly revealed no calcium dependence for SHIP1 (24). In general, the C2 domains of SHIP1/2 associate with anionic lipids in a calcium independent manner. To date, no SHIP1 separation of function mutants (e.g. point mutations) have been rationally designed to eliminate the reported interactions between the C2 domain and anionic lipids. Without definitive structural biochemistry data, the molecular basis of SHIP1 specificity for PS lipid binding outside of the catalytic phosphatase domain remains unclear.

Our HDX-MS experiments detected pY-mediated solvent exposure of residues in the CBL3 loop but only in the context of the dimeric SHIP1(ΔCTD) construct. Although the CBL3 motif might partially contribute to autoinhibition, charge reversal mutations in the CBL1 motif were sufficient to disrupt SHIP1 autoinhibition in vitro. Mutating the entire CBL1 motif (i.e. 7A mutant) also activates the minimal SHIP1 construct, mSHIP1(ΔCTD). By contrast, the charge neutralization mutations in the CBL1 motif (i.e. K741A/K743A/K747A) were unable to relieve autoinhibition. This could be due to the SH2 domain still having a sufficiently strong interaction with the C2 domain to mediate autoinhibition in the presence of the K741A/K743A/K747A mutations. There are several polar residues in the CBL1 motif that potentially hydrogen bond with the SH2 domain. Alternatively, there could be additional interactions between the SH2 domain and the CBL3 motif that facilitate autoinhibition in the context of the mSHIP1(ΔCTD) 3A mutant. Considering that the CBL1 and CBL3 motifs have opposite net charge, they could also interact with each other to form a surface that associates with the SH2 domain.

In vivo, alternative splicing of SHIP1/2 mRNA transcripts produce enzymes that lack the N-terminal SH2 domain (19, 41). Previous biochemical analysis indicates that these splice variants lack autoinhibition (24). The degree to which SHIP1 activity is modulated during cell signaling, however, will ultimately depend on cell type specific expression of receptors and peripheral membrane binding proteins that interact with either the N-or C-terminus of SHIP1. If cells contain an abundance of phosphotyrosine binding (PTB) or SH3 domain containing proteins that regulate SHIP1 localization via C-terminal domain interactions, it is expected that membrane localized SHIP1(ΔSH2) will exhibit more phosphatase activity. By contrast, cells that primarily rely on receptor tyrosine kinases (RTKs) to regulate SHIP1 plasma membrane localization will likely exhibit sustained PI3K signaling due to the reduced ability of a SHIP1(ΔSH2) splice variant to localize to the plasma membrane.

Overall, SHIP1 autoinhibition is regulated by weak intramolecular interactions between the N-terminal SH2 domain and the C2 domain. Although autoinhibitory interactions might restrict conformational changes in the phosphatase and C2 domains, our data suggests that SH2-mediated steric occlusion of the C2 domain functions to limit SHIP1 membrane association. As a result, SHIP1 activation is more dependent on tyrosine phosphorylated peptides which are produced at the plasma membrane during cell signaling. Our HDX-MS data and structure predictions also suggest that the SH2-C2 domain interaction does not directly block the active site. Although the linker connecting the N-terminal SH2 domains and central catalytic core could limit substrate accessibility to the active site, we did not observe a statistically significant change in solvent exposure across the linker sequence in the presence of pY peptides. Our results are more consistent with the N-terminal SH2 domain forming weak intramolecular interactions with the C2 domain that reduce the membrane binding frequency. Interactions with tyrosine phosphorylated peptides favor an open conformation of SHIP1. Dimerization of SHIP1 enhances membrane resident time by increasing the probability of the pY peptide engagement, resulting in high apparent phosphatase activity. Interaction with tyrosine phosphorylated peptides also increases the probability of collisions between anionic lipids and the PH-PP-C2 module, which promotes stable membrane docking and catalysis by SHIP1 (**Figure 7)**. This work provides a framework for future investigation of regulatory mechanisms, including small molecule drug design, that modulate SHIP1 localization and activity by strengthening or weakening the SH2-C2 interface.

**Figure 7.**
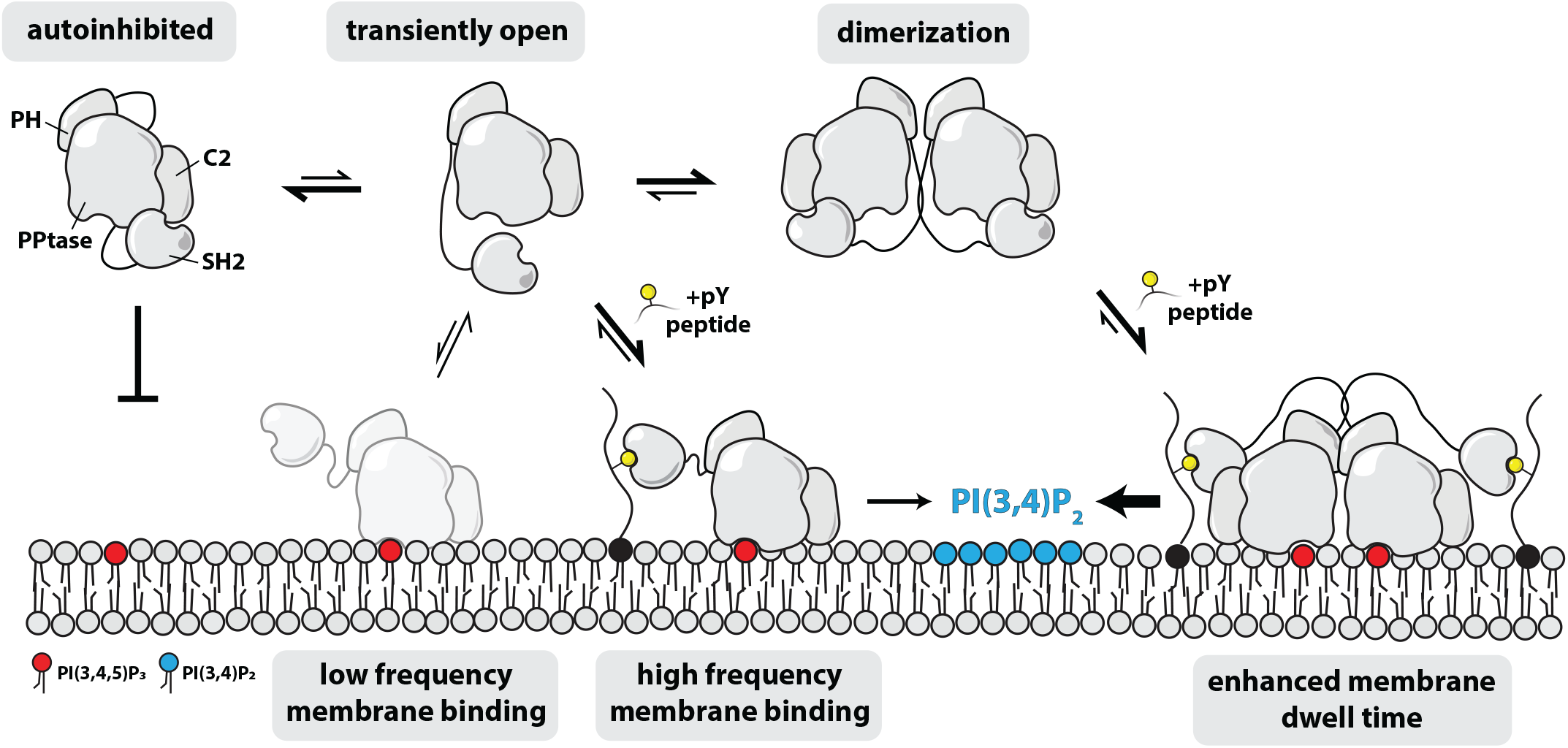
Mechanisms controlling SHIP1 membrane localization and phosphatase activity. In the autoinhibited state, the probability of SHIP1 membrane association is relatively low. Transient dissociation of the SH2-C2 domain interaction increases the probability of SHIP1 binding to anionic lipid. SHIP1 membrane interaction based solely on lipid interactions results in transient membrane dwell times. The presence of membrane tethered phosphotyrosine peptides enhances membrane localization and apparent phosphatase activity due to the increase in membrane resident time of SHIP1. Dimerization increases the membrane binding frequency and dwell time of SHIP1. This results in higher apparent phosphatase activity due to the increased membrane surface density observed in the presence of dimeric SHIP1.

## DATA AVAILABILITY

All the information needed for interpretation of the data is presented in the manuscript or the supporting information. Plasmids related to this work are available upon request.

## SUPPORTING INFORMATION

This article contains supporting information.

Supporting Information file – Figures, Tables, plasmid inventory, and peptide sequences

Figures S1-S6 Tables S1-S2

## AUTHOR CONTRIBUTIONS

Resources: E.E.D., R.K.T., S.D.H.

Experiments and investigation: E.E.D., H.G.N., M.A.P., R.K.T., J.E.B., S.D.H.

Data Analysis: E.E.D., H.G.N., M.A.P., J.E.B., S.D.H.

Conceptualization: E.E.D., J.E.B., S.D.H. Interpretation: E.E.D., H.G.N., M.A.P., J.E.B., S.D.H.

Data curation: E.E.D., H.G.N., M.A.P., S.D.H.

Writing – Review and editing: E.E.D., H.G.N., M.A.P., R.K.T., J.E.B., S.D.H.

Writing – Original draft: E.E.D. and S.D.H. Supervision: J.E.B. and S.D.H.

Project administration: S.D.H.

Funding acquisition: J.E.B. and S.D.H.

## MATERIALS & METHODS

### Molecular Biology

The plasmid containing the open reading frame for human Src-homology-2-domain-containing inositol 5-phosphatase 1 (SHIP1 or INPP5D) was purchased from Horizon Discovery (PerkinElmer), formerly known as Open Biosystems and Dharmacon. This SHIP1 clone represents isoform 2 (1-1189aa) and is missing V117 located in the disordered linker that follows the N-terminal SH2 domain (Uniprot Q92835, isoform 2). Gene fragments containing SHIP1 were cloned into FAST Bac1 or pETM vectors using Gibson Assembly (42). Each protein expression construct was screened for optimal yield and solubility using either bacteria (BL21 DE3 Star, Rosetta, etc.) or insect cells. Lentiviral packing vectors were purchased from Addgene. The psPAX2 plasmid was a gift from Didier Trono (Addgene plasmid #12260; http://n2t.net/addgene:12260; RRID:Addgene_12260). The pVSV-G plasmid was a gift from Akitsu Hotta (Addgene plasmid #138479) (43). All plasmids were sequenced by Azenta/Genewiz and Plasmidsaurus to ensure open reading frames lacked deleterious mutations. Refer to *Supporting Information* for complete list of plasmids used in this study.

### Protein Purification

#### mNG-mSHIP1(ΔCTD) (WT, 3D, 3A, 7A) and mNG-nSH2(PI3K)-SHIP1(PH-PP-C2)

The coding sequence of mSHIP1(ΔCTD) constructs was cloned into FASTBac1 vector with an N-terminal his6-TEV-mNG tag. Refer to *Supporting Information* for the exact amino acid sequences. BACMID DNA and baculovirus were generated as previously described (24, 33). High five insect cells were infected with baculovirus at 2% vol/vol and grown for 48 hours at 27ºC in ESF 921 Serum-Free insect cell culture medium (Expression Systems, Cat# 96-001-01). Cells were harvested by centrifugation at 3500 rpm for 10 minutes and transferred into 50 mL conicals using 1x PBS [pH 7.2] to resuspend and wash the cells. The cells were pelleted and resuspended into an equal volume of 1x PBS [pH 7.2] containing 10% Glycerol and 2x protease inhibitors (1 sigma protease inhibitor tablet per 50 mL of buffer). The pellets were frozen down in a -80 ºC until purification. To purify SHIP1 proteins, insect cell pellets were thawed in room temperature water and resuspended in buffer containing 30 mM Tris [pH 8.0], 10 mM imidazole, 5% Glycerol, 400 mM NaCl, 1 mM PMSF, 2 mM BME, 1 Sigma protease inhibitor tablet per 100 mL of buffer, and 100 µg/mL DNase. Cells were lysed using dounce homogenizer. The lysate was centrifuged at 35,000 rpm at 4ºC in a Beckman Ti45 rotor. High speed supernatant (HSS) was pooled and batch bound to 5-10 mL of Ni-NTA agarose resin (Qiagen, Cat# 30230) at 4ºC for 2 hours stirring in a beaker. The HSS and resin was poured into 50 mL conical tubes, centrifuged, and then washed with buffer containing 20 mM Tris [pH 8.0], 30 mM imidazole, 400 mM NaCl, 5% glycerol, and 2 mM BME. The resin bound to his6-mNG-mSHIP1(ΔCTD) constructs was transferred to a gravity flow column and washed with at least 100 mL of additional wash buffer. The protein was then eluted with buffer containing 20 mM Tris [pH 8.0], 300 mM imidazole, 400 mM NaCl, 5% glycerol, and 2 mM BME. Peak fractions were pooled, and his6-TEV protease (poly-E tail) was added to the elution to cleave off the poly histidine tag. SHIP1 proteins were dialyzed overnight into 4L reservoir of buffer containing 20 mM Tris [pH 8.0], 150 mM NaCl, and 2 mM BME to reduce the salt concentration for ion exchange chromatography. The next day, dialysate was transferred to a 50 mL tube and centrifuged at 3500 rpm at 4 ºC to clarify. The supernatant was passed through a 0.22 µm syringe filter to remove any precipitation. Buffer containing 20 mM Tris [pH 8.0], 1 mM TCEP is added to the dialysate in a 1:2 ratio to lower the salt concentration to 100 mM NaCl. The protein was then bound to a 1 mL HiTrap CaptoQ anion exchange column (Cytiva Cat# 11-0013-02) equilibrated in 20 mM Tris [pH 8.0], 100 mM NaCl, 1 mM TCEP. Proteins were eluted with a linear salt gradient from 0.1-1M NaCl over 22.5 minutes at a flow rate of 1.5mL/min. The mNG-mSHIP1(ΔCTD) constructs elute at 300-450 mM NaCl. Peak fractions were pooled and concentrated in 100 kDa MWCO Amicon Ultra-4 spin concentrator (Sigma Millipore, Cat#: UFC810024) to 0.5-1mL. The protein was then loaded onto a 24 mL Superdex200 10/300 GL (GE Healthcare, Cat# 17-5174-01) size exclusion chromatography column equilibrated in buffer containing 20 mM Tris [pH 8.0], 150-200mM NaCl, 10% Glycerol, and 1 mM TCEP. The peak fractions were pooled and concentrated using a 100 kDa MWCO Amicon Ultra-4 spin concentrator. Proteins were aliquoted and flash frozen at concentration of 1-30 µM using liquid nitrogen. The purification protocol for mSHIP1(ΔCTD), wild-type and mutants, was identical. Biophysical parameters of mNG-mSHIP1(ΔCTD): pI = 6.37, ε_280_ = 131100 M^-^1•cm^-1^, 112.9 kDa. Biophysical parameters for mNG-PI3K(nSH2)-SHIP1(PH-PP-C2): pI = 6.17, ε_280_ = 138090 M^-^1•cm^-1^, 115 kDa.

#### SHIP1 (full-length and ΔCT)

The coding sequence of full-length human SHIP1 (1-1189 aa) was cloned into a modified FastBac1 vector containing an N-terminal his6-TEV or his6-TEV-mNG fusion. High five insect cells were infected with baculovirus using an optimized multiplicity of infection (MOI), typically 2% vol/vol, which we empirically determined based on small-scale test expressions (25-50 mL culture). Infected cells were typically grown for 48 hours at 27ºC in ESF 921 Serum-Free insect cell culture medium (Expression Systems, Cat# 96-001-01). Cells were harvested by centrifugation, transferred to 50 mL tubes by washing with 1x PBS [pH 7.2], pelleted and resuspended in 10 mL of 1x PBS [pH 7.2], 10% Glycerol, and 2x protease inhibitor cocktail (1 Sigma protease inhibitor tablet per 50 mL of buffer) and then stored in the -80ºC freezer. For purification, frozen cell pellets were thawed in an ambient water bath and lysed into buffer containing 30 mM Tris [pH 8.0], 10 mM imidazole, 5% glycerol, 400 mM NaCl, 1 mM PMSF, 2 mM BME, Sigma protease inhibitor cocktail EDTA free per 100 mL lysis buffer, and 100 µg/mL DNase using a dounce homogenizer. Lysate was then centrifuged at 35,000 rpm for 60 minutes in a Beckman Ti 45 (Cat#: 339160) rotor at 4ºC. High speed supernatant (HSS) was then batch bound to 5 mL of Ni-NTA Agarose (Qiagen, Cat# 30230) resin at 4ºC for 2 hours stirring in a beaker. The resin and HSS was collected in 50 mL tubes, centrifuged, and washed with buffer containing 20 mM Tris [pH 8.0], 30 mM imidazole, 300-400 mM NaCl, 5% glycerol, and 2 mM BME before being transferred to gravity flow column. NiNTA resin with bound his6-TEV-mNG-SHIP1 was then washed with an additional 100 mL of 20 mM Tris [pH 8.0], 30 mM imidazole, 300-400 mM NaCl, 5% glycerol, and 2 mM BME buffer and then eluted into buffer containing 300 mM imidazole. Peak fractions were pooled and desalted using a G25 Sephadex desalting column (GE Healthcare/Cytiva, Cat#: 17-5087-01) equilibrated in 20 mM Tris [pH 8.0], 100 mM NaCl, and 1mM TCEP buffer. Any precipitation was removed via 0.22 µm syringe filtration. Clarified SHIP1 fractions were then bound to a HiTrap 1 mL CaptoQ anion exchange column (Cytiva, cat#11-0013-02) equilibrated in 20 mM Tris [pH 8.0], 100 mM NaCl, and 1 mM TCEP buffer. Proteins were resolved over a 10-100% linear gradient (0.1-1 M NaCl, 30 minutes, 1.5 mL/min flow rate). His6-TEV-mNG-SHIP1 elutes in the presence of 200-400 mM NaCl. Peak fractions containing SHIP1 were pooled, incubated with 400 μL of 243 μM his6-TEV protease (S291V) with poly-R tail for ∼12-16 hours at 40C, concentrated in a 100 kDa MWCO Amicon concentrator (Sigma Millipore,), and then loaded onto a 24 mL Superdex 200 10/300 GL (GE Healthcare, Cat# 17-5174-01) size exclusion column equilibrated in 20 mM Tris pH [8.0] 200 mM NaCl, 10% glycerol, 1 mM TCEP, and 0.05% CHAPS buffer (SHIP1 (ΔCT) final buffer contained no CHAPS and 0.5 mM TCEP). Peak fractions were concentrated in a 100 kDa MWCO Amicon Ultra-4 spin concentrator and flash frozen at a final concentration of 1-30 µM using liquid nitrogen. Purification schemes for full-length SHIP1, SHIP1(ΔSH2), and SHIP1(ΔCT) were identical. Biophysical parameters of FL SHIP1: pI = 7.37, ε_280_ = 106,690 M^-1^•cm^-1^, 133.3 kDa.

#### mNG-SHIP1(PH-PP-C2)

BL21 (DE3) Star bacteria were transformed with his10-TEV-mNG-GGGGG-SHIP1 (mNG-PH-PP-C2, 191-878aa), wild-type and mutant (K741D/K743D/K747D). and plated on LB agar containing 50 μg/ml kanamycin. The following day, 50mL of TPM media (20g tryptone, 15g yeast extract, 8g NaCl, 2g Na_2_HPO_4_, and 1g KH_2_PO_4_ per liter) containing 50 μg/ml kanamycin was inoculated with one bacterial colony from the agar plate. This culture was grown for 12-16 hours at 27ºC to an OD600 = 2–3, before being diluted to an OD_600nm_ of 0.05 in 2 liters of TPM media. These cultures were grown at 37ºC to an OD_600nm_ of 0.8, shifted to 30ºC for 1 hour, and then bacteria were induced with 50 μM isopropyl β-d-1-thiogalactopyranoside (IPTG) to express mNG-SHIP1(PH-PP-C2). After 6-8 hours of expression at 30°C, bacterial cultures were harvested by centrifugation, transferred to 50 mL tubes by washing with 1x PBS [pH 7.2], pelleted, and then stored in the -80°C freezer. The PH-PP-C2 (K741D/K743D/K747D) protein was diluted into 2 liters of TB media and expressed at 18 ºC for 18-20 hourse before harvesting. For purification, frozen cell pellets were thawed in an ambient water bath and lysed into buffer containing 50 mM Na_2_HPO_4_ [pH 8.0], 400 mM NaCl, 1 mM PMSF, 0.4 mM BME, and 100 µg/mL DNase by micro-tip sonication (12-24 x 5 second pulses, 40% amplitude). Lysate was then centrifuged at 15,000 rpm (35,000xg) for 60 minutes in a Beckman JA-20 rotor at 4ºC. During lysate centrifugation, a 5mL HiTrap chelating column (Cat#) was charged with 50 mL of 100mM CoCl_2_, generously washed with water (phosphate buffer will precipitate unchelated CoCl_2_ if not thoroughly removed) and equilibrated with lysis buffer lacking PMSF and DNase (>0.4mM BME will destroy charged HiTrap column and turn column brown or black). High speed supernatant (HSS) was then circulated over charged 5mL HiTrap column for 2 hours. Captured protein was washed with 100mL of 50 mM Na_2_HPO_4_ [pH 8.0], 400 mM NaCl, 10mM imidazole, and 0.4 mM BME and then eluted into buffer containing 500 mM imidazole. Peak fractions containing SHIP1 were pooled with 400 μL of 243 μM his6-TEV protease (S291V) with poly-R tail and dialyzed in 4L of 25 mM Na_2_HPO_4_ [pH 8.0], 400 mM NaCl, and 0.4 mM BME buffer at 4°C for ∼12-16 hours to remove imidazole. The next day, the charged HiTrap column was equilibrated with dialysis buffer and dialyzed protein was recirculated for 2 hours at 4°C to remove his6-TEV protease and uncleaved his10-TEV-mNG-GGGGG-SHIP1(PH-PP-C2). Unbound protein was then concentrated in a 30 kDa MWCO Amicon concentrator (Sigma Millipore) and then loaded onto a 24 mL Superdex 200 10/300 GL (GE Healthcare, Cat# 17-5174-01) size exclusion column equilibrated in 20 mM Tris pH [8.0], 200 mM NaCl, 10% glycerol, and 1 mM TCEP buffer. Peak fractions were concentrated in a 30 kDa MWCO Vivaspin 6 centrifuge tube and flash frozen at a final concentration of 10-30 µM using liquid nitrogen. Biophysical parameters of mNG-SHIP1(PH-PP-C2): pI = 7.43, ε_280_ = 116,660 M^-1^•cm^-1^, 94.4 kDa.

#### mNG-SHIP1 (RBD)

BL21 (DE3) Star bacteria were transformed with his10-mNG-GGGGG-SHIP1 (RBD,151-290aa) and plated onto an LB agar plate containing 50 μg/ml kanamycin. To express the protein, 50 mL of TPM media with 50 μg/ml kanamycin was inoculated with one bacterial colony and grown at 27ºC for 12-16 hours until OD600 = 1.5-2. The culture was the diluted into 2L of TB media to an OD600 = 0.05. The cultures were grown at 37ºC for 1.5-3 hours until OD600 = 0.6-0.8. The temperature was shifted to 18 ºC for 1 hour before inducing expression of mNG-SHIP1 (RBD) with 100 μM isopropyl β-d-1-thiogalactopyranoside (IPTG). The cultures were then allowed to grow 16-20 hours before harvesting. The bacterial cultures were harvested by centrifugation, and the pellets were resuspended in 1x PBS [pH 7.2] and transferred to 50 mL conicals for storage at -80ºC. Frozen cell pellets were thawed for purification in an ambient water bath and lysed into buffer containing 50 mM Na_2_HPO_4_ [pH 8.0], 400 mM NaCl, 1 mM PMSF, 0.4 mM BME, and 100 µg/mL DNase. The cells were then lysed by sonication (24 pulses each 5 seconds at 40% amplitude). The lysate was then centrifuged at 15,000 rpm at 4 ºC for 1 hour in JA-20 rotor (Beckman). The high-speed supernatant was then circulated for 2 hours over a HiTrap chelating column loaded with 50 mL 200mM CoCl_2_ that was equilibrated in lysis buffer excluding PMSF and DNase. After circulation, the HiTrap column was washed with 100 mL of wash buffer containing 50 mM Na_2_HPO_4_ [pH 8.0], 400 mM NaCl, 10mM imidazole, and 0.4 mM BME and then eluted into buffer with 500 mM imidazole. The peak fractions were pooled and 400 uL of his6-TEV protease was added to cleave the histidine tag and dialyzed into 4L of 25 mM Na_2_HPO_4_ [pH 8.0], 400 mM NaCl, and 0.4 mM BME buffer overnight at 4ºC. The following morning, the dialysate was recirculated over the HiTrap column to trap the cleaved his tag and the TEV protease. The dialysate was then concentrated to ∼0.5 mL in 30 kDa MWCO Amicon concentrator (Sigma Millipore) to load on the Superdex200 10/300 GL column (GE Healthcare, Cat# 17-5174-01) equilibrated in buffer containing 20 mM HEPES [pH 7], 200 mM NaCl, 10% glycerol, 1 mM TCEP. Peak fractions were determined by the A280 and SDS-PAGE and the fractions were pooled and concentrated using a 30 kDa MWCO Amicon concentrator (Sigma Millipore). The concentrated protein was aliquoted and flash frozen at a concentrated at 70-180 μM with liquid nitrogen and subsequently stored at -80ºC. Biophysical parameters of mNG-SHIP1(RBD): pI = 6.45, ε_280_ = 47,330 M^-1^•cm^-1^, and 42.7 kDa for the monomeric species.

#### Btk-SNAP

The PI(3,4,5)P_3_ binding domain derived from Btk was recombinantly expressed and purified as a his6-SUMO-Btk(PH-TH,R49S/K52S)-SNAP fusion protein using a protocol previously described (26). This construct contains mutations (i.e. R49S/K52S) in the peripheral lipid binding domain, which eliminate dimerization and the valency of PI(3,4,5)P_3_ lipid interactions (Chung et al. 2019; Wang et al. 2015). The his6-SUMO-Btk(PH-TH,R49S/K52S)-SNAP construct was recombinantly expressed in BL21 Star *E. coli*. Bacteria were initially grown at 37°C in Terrific Broth to an OD600 of 0.8. Cultures were then shifted to 18°C for 1 hr, induced with 0.1 mM IPTG, and allowed to express protein for 20 hr at 18°C before being harvested. Cells were lysed into 50 mM NaPO_4_ (pH 8.0), 400 mM NaCl, 0.5 mM BME, 10 mM Imidazole, and 5% glycerol. Lysate was then centrifuged at 16,000 rpm (35,172 × *g*) for 60 min in a Beckman JA-20 rotor chilled to 4°C. Lysate was circulated over 5 mL HiTrap Chelating column (GE Healthcare, Cat# 17-0409-01), wash, and then eluted with linear gradient of lysis buffer containing 0-500 mM imidazole (8 CV, 40 mL total, 2 mL/min flow rate). Peak fractions were pooled and combined with SUMO protease Ulp1 to cleave the his6-SUMO tag. The protein was dialyzed against 4 L of buffer containing 20 mM Tris [pH 8.0], 200 mM NaCl, and 0.5 mM BME for 18 hr at 4°C. The next day, cleaved Btk was recirculated for 1 hr over a 5 mL HiTrap Chelating column. Flow-through containing Btk-SNAP was then concentrated in a 5 kDa MWCO Vivaspin 20 before being loaded on a Superdex 75 size-exclusion column equilibrated in 20 mM Tris [pH 8.0], 200 mM NaCl, 10% glycerol, 1 mM TCEP. Peak fractions containing Btk-SNAP were pooled and concentrated to a concentration of 30 µM before snap-freezing with liquid nitrogen and storage at –80°C. For labeling, Btk-SNAP was combined with a 1.5× molar excess of SNAP-Surface Alexa546 dye (NEB, Cat# S9132S) and incubated overnight at 4ºC. The next day, Btk-SNAP-AF546 was desalted into buffer containing 20 mM Tris [pH 8.0], 200 mM NaCl, 10% glycerol, 1 mM TCEP using a PD10 column. The protein was then concentrated using a Amicon filter and loaded onto a Superdex 75 column separate Btk-SNAP-AF546 from free AF546 dye. The peak elution was pooled, concentrated, aliquoted, and flash frozen with liquid nitrogen.

### Preparation of supported lipid bilayers

Supported lipid bilayers (SLBs) were created to maximize the membrane fluidity at room temperature (i.e. 23ºC). The membrane composition is primarily DOPC (i.e. 94-98%), which has a melting temperature of -20ºC (44). Unless otherwise noted, SLBs also include PI(3,4,5)P_3_ as the substrate for SHIP1 and 20% phosphatidylserine to make the membrane more electronegative like the plasma membrane. In addition, some experiments include maleimide functionalized lipids in order conjugate pY peptides to the membrane. Although the SLBs lack complexity that exists on the plasma membrane in vivo, this experimental platform enables the SHIP1 phosphatase activity and membrane binding interactions to analyze using a reductionist biochemistry approach.

The following lipids were used to generated small unilamellar vesicles (SUVs): 1,2-dioleoyl-sn-glycero-3-phosphocholine (18:1 DOPC, Avanti #850375C), 1,2-dioleoyl-*sn-*glycero-3-phospho-L-serine (18:1 DOPS, Avanti #840035C), 1,2-dioleoyl-sn-glycero-3-phosphoethanolamine-N-[4-(p-maleimidomethyl)cyclohexane-carboxamide] (18:1 MCC-PE, Avanti #780201C), and D-myo-phosphatidylinositol 3,4,5-trisphosphate (PI(3,4,5) P_3_ diC16, Echelon P-3916-100ug). To prepare of small unilamellar vesicles, 2 µmoles total lipids were combined in a 35 mL glass round bottom flask containing 2 mL of chloroform. Lipids were dried to a thin film using rotary evaporation with the glass round-bottom flask submerged in a 42ºC water bath. After evaporating all the chloroform, the round bottom flask was place under a vacuum for 15 minutes. The lipid film was then resuspended in 2 mL of PBS [pH 7.2], making a final concentration of 1 mM total lipids. All lipid mixtures expressed as percentages (e.g. 98% DOPC, 2% PI(3,4,5)P_3_) are equivalent to molar fractions. For example, a 1 mM lipid mixture containing 98% DOPC and 2% PI(3,4,5)P_3_ is equivalent to 0.98 mM DOPC and 0.02 mM PI(3,4,5)P_3_. To generate 50 nm SUVs, 1 mM total lipid mixtures were extruded through a 0.05 µm pore size 19 mm polycarbonate (PC) membrane (Cytiva Whatman, Cat# 800308) with filter supports (Cytiva Whatman, Cat# 230300) on both sides of the PC membrane.

Supported lipid bilayers were formed on 25x75 mm coverglass (IBIDI, #10812) as previously described (33). In brief, coverglass was first cleaned with 2% Hellmanex III (ThermoFisher, Cat#14-385-864) and then etched with Pirahna solution (1:3, hydrogen peroxide:sulfuric acid) for 15 minutes. Etched coverglass, washed extensively with MilliQ water again, is rapidly dried with nitrogen gas before adhering to a 6-well sticky-side chamber (IBIDI, Cat# 80608). SLBs were formed by flowing 0.25 mM total lipid concentration of 50 nm SUVs diluted in 1x PBS pH [7.2] into an assembled IBIDI chamber. SUVs were incubated in the IBIDI chamber for 30 minutes and then washed with 4 mL of PBS [pH 7.4] to remove non-absorbed SUVs. Membrane defects are blocked for 5 minutes with a solution of clarified 1 mg/mL beta casein (ThermoFisher, Cat# 37528) in 1x PBS [pH 7.2]. After blocking membranes with 1 mg/mL beta casein for 5 minutes, bilayers were washed with 4 mL of 1x PBS. Reaction/imaging buffer was exchanged into the chamber prior to each experiment.

Conjugation of pY peptides to supported membranes was achieved as previously described (24). Supported membrane containing 2% MCC-PE lipids were used to covalently couple phosphorylated peptides (**<u>C</u>**KTEAENTIT(pY)SLIK, Elim Biopharmaceuticals) derived from the ITIM sequence FcγRIIB (45). Phosphorylated peptides correspond to the A cysteine (bold) was added to the N-terminus of the peptide to form a covalent interaction with the maleimide lipid headgroup of the MCC-PE lipids. For membrane coupling, 100 µL of 10 µM pY-ITIM peptides diluted in a 1x PBS [pH 7.2] and 0.1 mM TCEP buffer was added to the IBIDI chamber and incubated for 2 hours at 23ºC. Unreacted MCC-PE lipids were then quench 15 minutes using 1x PBS [pH 7.4] with 5 mM beta-mercaptoethanol (BME). Quenched membranes were then washed and stored in 1x PBS [pH 7.4].

### Lipid phosphatase activity measurements

All 5-phosphatase activity measurements were performed using SHIP1 proteins with N-terminal mNeonGreen (mNG) fusions. For simplicity, the mNG notation was omitted from all the graphs shown in Figures 1, 4, 5, S5. The kinetics of SHIP1 mediated dephosphorylation PI(3,4,5)P_3_ to generate PI(3,4)P_2_ was monitored on supported lipid bilayers formed on piranha etched glass in an IBIDI chamber using TIRF microscopy. Phosphatase reactions contained 20 mM HEPES [pH 7.0], 100 mM NaCl, 1 mM ATP, 5 mM MgCl_2_, 0.5 mM EGTA, 200 µg/mL beta casein, 20 mM BME, 20 mM glucose, and 0.32 mg/mL glucose oxidase, 50 µg/mL catalase, and 1 mM Trolox. Membranes contained the following initial composition: 98% DOPC, 2% PI(3,4,5)P_3_, or 96% DOPC, 2% PI(3,4,5) P_3_, 2% MCC-PE or 78% DOPC, 20% DOPS, 2% PI(3,4,5)P_3_. Membrane compositions are indicated in the figure legends. All kinetic measurements utilized 20 nM Btk-SNAP-AF546 (PH domain; PI(3,4,5)P_3_ sensor) to visualize the dephosphorylation of PI(3,4,5)P_3_ by TIRF microscopy. Assuming a footprint of 0.72 nm^2^ for DOPC lipids, we calculated a density of 27778 lipids/µmr_2_ for 2% PI(3,4,5)P_3_ (46, 47).

### Oxygen scavenging system

Glucose oxidase (32 mg/mL, 100x stock) and catalase (5 mg/mL, 100x stock) were solubilized in 20 mM HEPES [pH 7.0], 150 mM NaCl, 10% glycerol, and 1 mM TCEP buffer, and then flash frozen in liquid nitrogen and stored at -80°C. Trolox (200mM, 100x stock) is UV treated and stored at -20°C. Approximately 10 minutes before imaging, 100x oxygen scavenger stocks were diluted in kinase buffer containing enzymes/biosensors to achieve a final concentration of 320 µg/mL glucose oxidase (Serva, #22780.01 *Aspergillus niger*), 50 µg/mL catalase (Sigma, #C40-100MG Bovine Liver), and 2 mM Trolox (Cayman Chemicals, cat #10011659). Trolox was prepared using a previously described protocol that utilized UV irradiation to drive the formation of a quinone species (48, 49).

### HDX-MS sample preparation

HDX reactions comparing mNG-SHIP1(ΔCTD) +/-40 µM pY peptide were carried out in a 10 µl reaction volume containing 15 pmol (1.5µM) of mNG-SHIP1(ΔCTD). The exchange reactions were initiated by the addition of 7.5 µL of D_2_O buffer (20 mM HEPES pH 7.5, 200 mM NaCl, 0.5 mM TCEP, 94.25% D_2_O [v/v]) to 2.5 µL of protein (final D_2_O concentration of 70.7%). Reactions proceeded for 3s, 30s, 300s and 3000s at 20°C before being quenched with ice cold acidic quench buffer, resulting in a final concentration of 0.6 M guanidine HCl and 0.9% formic acid post quench. HDX reactions comparing apo mNG-mSHIP1(ΔCTD) +/-40 µM pY peptide were carried out in 10 µL reaction volumes containing 15 pmol of protein. The exchange reactions were initiated by the addition of 7 µL of D_2_O buffer (20 mM HEPES pH 7.5, 50 mM NaCl, 95.24% D_2_O [v/v]) to 3 µL of protein (final D_2_O concentration of 66.67% [v/v], 1.5 µM mNG-mSHIP1(ΔCTD), 40 µM pY peptide). Reactions proceeded for 3s, or 300s before being quenched with ice cold acidic quench buffer, resulting in a final concentration of 0.6 M guanidine HCl and 0.9% formic acid post quench. All conditions and timepoints were created and run in independent triplicate. Samples were flash frozen immediately after quenching and stored at -80°C until injected onto the ultra-performance liquid chromatography (UPLC) system for proteolytic cleavage, peptide separation, and injection onto a QTOF for mass analysis, described below.

### Protein Digestion and MS/MS Data Collection

Protein samples involving mNG-SHIP1(ΔCTD) were rapidly thawed and injected onto an integrated fluidics system containing a HDx-3 PAL liquid handling robot and climate-controlled (2°C) chromatography system (LEAP Technologies), a Waters Acquity UPLC I-Class Series System, as well as an Impact HD QTOF Mass spectrometer (Bruker). mNG-mSHIP1(ΔCTD) protein samples were run using a Dionex Ultimate 3000 UHPLC system. The full details of the automated LC systems were previously described in (50). mNG-SHIP1(ΔCTD) samples were run over an immobilized pepsin column (Affipro; AP-PC-001) at 200 µL/min for 4 minutes at 2°C. The resulting peptides were collected and desalted on a C18 trap column (Acquity UPLC BEH C18 1.7 µm column (2.1 × 5 mm); Waters 186004629). The trap was subsequently eluted in line with an ACQUITY 300Å, 1.7 μm particle, 100 × 2.1 mm BEH C18 UPLC column (Waters), using a gradient of 3-10% B (Buffer A 0.1% formic acid; Buffer B 100% acetonitrile) over 1.5 minutes, followed by a gradient of 10-25% B over 4.5 minutes, followed by a gradient of 25-35% B over 5 minutes, finally after 1 minute at 35% B a gradient of 35-80% B over 1 minute was used. mNG-mSHIP1(ΔCTD) samples were run over one immobilized pepsin column (Trajan; ProDx protease column, 2.1 mm x 30 mm PDX.PP01-F32) at 200 µL/min for 3 minutes at 8°C. The resulting peptides were collected and desalted on a C18 trap column (Acquity UPLC BEH C18 1.7µm column (2.1 x 5 mm); Waters 186003975). The trap was subsequently eluted in line with an ACQUITY 1.7 μm particle, 100 × 1 mm2 C18 UPLC column (Waters), using a gradient of 3-35% B (Buffer A 0.1% formic acid; Buffer B 100% acetonitrile) over 11 minutes immediately followed by a gradient of 35-80% over 5 minutes. Mass spectrometry experiments acquired over a mass range from 150 to 2200 m/z using an electrospray ionization source operated at a temperature of 200°C and a spray voltage of 4.5 kV.

### Peptide identification

Peptides were identified from the non-deuterated samples of mNG-SHIP1(ΔCTD) using data-dependent acquisition following tandem MS/MS experiments (0.5 s precursor scan from 150-2000 m/z; twelve 0.25 s fragment scans from 150-2000 m/z). MS/MS datasets were analysed using FragPipe v18.0 and peptide identification was carried out by using a false discovery-based approach using a database of purified proteins and known contaminants (51–53). MSFragger was utilized, and the precursor mass tolerance error was set to -20 to 20ppm. The fragment mass tolerance was set at 20ppm. Protein digestion was set as nonspecific, searching between lengths of 4 and 50 aa, with a mass range of 400 to 5000 Da. mNG-mSHIP1(ΔCTD) MS/MS datasets were analysed using PEAKS7 (PEAKS), and peptide identification was carried out by using a false discovery-based approach, with a threshold set to 0.1% using a database of purified proteins and known contaminants. The search parameters were set with a precursor tolerance of 20 ppm, fragment mass error 0.02 Da, charge states from 1-8, leading to a selection criterion of peptides that had a -10logP score of 32.6.

### Mass Analysis of Peptide Centroids and Measurement of Deuterium Incorporation

HD-Examiner Software (Sierra Analytics) was used to automatically calculate the level of deuterium incorporation into each peptide. All peptides were manually inspected for correct charge state, correct retention time, appropriate selection of isotopic distribution, etc. Deuteration levels were calculated using the centroid of the experimental isotope clusters. Results are presented as relative levels of deuterium incorporation and the only control for back exchange was the level of deuterium present in the buffer (70.7% for mNG-SHIP1(ΔCTD) and 66.67% for mNG-mSHIP1(ΔCTD). Differences in exchange in a peptide were considered significant if they met all three of the following criteria: ≥5% change in exchange, ≥0.4 Da difference in exchange, and a p value <0.01 using a two tailed student t-test for mNG-SHIP1(ΔCTD) experiment or ≥5% change in exchange, ≥0.4 Da difference in exchange, and a p value <0.01 using a two tailed student t-test for mNG-mSHIP1(ΔCTD) experiment. Slight differences in significance thresholds between experiments is due to small differences in the system used for analysis. Samples were only compared within a single experiment and were never compared to experiments completed at a different time with a different final D_2_O level. The data analysis statistics for all HDX-MS experiments are in **Table S1** and **Table S2** according to published guidelines (54). The mass spectrometry proteomics data have been deposited to the ProteomeXchange Consortium via the PRIDE partner repository (55) with the dataset identifier **PXD058704** for mNG-SHIP1(ΔCTD) data and **PXD061719** for mNG-mSHIP1(ΔCTD) data.

### Cell culture

PLB-985 neutrophil-like cells were grown in suspension in RPMI 1640 + GlutaMAX media containing 25 mM HEPES (Life Technologies, cat #72400047), 9% fetal bovine serum (FBS), penicillin (100 units/ml) (Life Technologies, cat #15140122), streptomycin (100 µg/ml) (Life Technologies, cat #15140122). These cells were provided by Professor Sean Collins (University of California at Davis) and were validated by RNA-seq compared to primary human neutrophils (56). Cell lines were stored in a humidified incubators at 37°C in the presence of 5% CO2 and split three times per week to maintain densities between 0.1-2 x 10^6^ cells/mL. PLB-985 cells were differentiated into a neutrophil-like state by culturing 0.2 x 10^6^ cells/mL for 6 to 7 days in RPMI media supplemented with 2% FBS, penicillin (100 units/ml), streptomycin (100 µg/ml), 1.3% DMSO, and 2% Nutridoma-CS (Sigma, cat #11363743001). Nutridoma-CS was added to increase the chemotactic response of the cells.

Human embryonic kidney (HEK) 293T Lenti-X were obtained from Takara (cat# 632180). These cells transformed with adenovirus type 5 DNA and expresses the SV40 large T antigen. HEK293T Lenti-X cells were cultured in DMEM + GlutMAX + High Glucose (4.5 g/L) + sodium pyruvate (110 mg/L) (Life Technologies, cat #10569010) supplemented with 10% FBS (Sigma, cat# F4135-500ML), penicillin (100 units/ml), and streptomycin (100 µg/ml). Cells were grown in 10 cm dishes in humidified incubators at 37°C in the presence of 5% CO_2_ and split at a confluency of 80-90% every 2-3 days. HEK293T Lenti-X cells were split using using 1.5mL of Cellstripper (Corning, cat# 25-056-Cl). Cellstripper was then quenched with 8.5 mL complete DMEM media containing 10% FBS. Cells were diluted 1:10 and seeded on a new 10cm dish containing a total volume of 10 mL complete DMEM media warmed to 37°C.

### Lentivirus production

Lentivirus was generated by transfecting 60-70% confluent HEK293 Lenti-X cells in a 10-cm plate with 6.7 µg psPAX2 (2^nd^ generation lentiviral packaging plasmid, Addgene #12260), 0.85 µg pVSV-G (Expresses VSV-G envelop protein for pseudotyping NanoMEDIC particle, Addgene #138479) (43), 7.5 µg of transfer lentiviral vector containing gene of interest, and 30 µL of 1 mg/mL polyethyleneimine (PEI) in 0.5 mL Opti-Mem (ThermoFisher, cat#31985070). This mixture was incubated for 15 minutes at room temperature before adding dropwise to plated HEK293 Lenti-X cells. Transfection was performed in complete media and media was not changed at any point during the transection or lentiviral production process. Media containing lentivirus was harvested 48 hours after beginning the transfection. Media containing lentivirus was clarified by centrifugation and then concentrated using Lenti-X concentrator (Takara, cat# 631231). Precipitated virus was resuspended in 0.4 mL of complete RPMI media (9% FBS) and then stored at -80°C. To transduce PLB-985 cells, 0.4 mL of concentrated lentivirus was added with 5 mL of undifferentiated cells at a density of 0.2 x 10^6^ cells/mL containing a final concentration of 8 µg/mL polybrene (Millipore, Cat# TR-1003-G, 10 mg/mL stock, 1250x) in the cell culture media. To increase transduction efficiency, spinoculation was performed in a 6-well dish spun at 1000 x g for 1 hour at 33ºC. Infected PLB-985 cells were passaged at least one time before differentiating into the neutrophil-like state (see *Cell Culture* for protocol).

### Live cell imaging

To image cells, 25×75 mm coverslips were cleaned overnight with 2% Hellmanex III (ThermoFisher, Cat# 14-385-864). The coverslips were rinsed well with MilliQ water and dried with N_2_ gas and attached to a 6-well IBIDI flow cell chamber (IBIDI sticky-Slide VI 0.4, cat #80608).

Cells were imaged in Hank’s balance salt solution (HBSS) containing 20 mM HEPES [pH 7.2], 150 mM NaCl, 4 mM KCl, 1 mM MgCl_2_, and 10 mM glucose that was warmed to 37°C. A 10 µg/mL fibronectin (Sigma, cat# F1141, 1 mg/mL stock concentration) solution diluted in 1x PBS was added to each well of the IBIDI chamber and incubated for 60 minutes at room temperature. Unbound fibronectin was washed out with 1x PBS and then blocked with 0.2% BSA (bovine serum albumin fatty/endotoxin free, Sigma, cat# B4501) dissolved in HBSS for 5 minutes. Differentiated PLB-985 (6 days post differentiation) were prepared for imaging by centrifuging 500 µL of cells at 100 x *g* for 10 minutes. The RPMI media was aspirated off and the cells were resuspended in warm HBSS. The cells were flowed into the IBIDI chamber and allowed to adhere for 15-20 minutes. The cells were imaged under a Okolab heated stage (Cat#: H301-K-FRAME) warmed to 37°C.

### Microscope hardware and imaging acquisition

Membrane binding and lipid dephosphorylation reactions reconstituted on supported lipid bilayers (SLBs) were visualized using an inverted Nikon Eclipse Ti2 microscope using a 100x Nikon (1.49 NA) oil immersion TIRF objective. TIRF-M images of SLBs were acquired using an iXion Life 897 EMCCD camera (Andor Technology Ltd., UK). Fluorescently labeled proteins were excited with either a 488 nm, 561 nm, or 637 nm diode laser (OBIS laser diode, Coherent Inc. Santa Clara, CA) controlled with a Vortran laser drive with acousto-optic tunable filters (AOTF) control. The power output measured through the objective for single particle imaging was 1-2 mW. Excitation light was passed through the following dichroic filter cubes before illuminating the sample: (1) ZT488/647rpc and (2) ZT561rdc (ET575LP) (Semrock). Fluorescence emission was detected on the iXion Life 897 EMCCD camera position after a Nikon emission filter wheel housing the following emission filters: ET525/50M, ET600/50M, ET700/75M (Semrock). All in vitro biochemistry experiments were reconstituted at room temperature (23ºC), while live cell imaging was performed at 37ºC using an Okolab heated stage. Microscope hardware was controlled by Nikon NIS elements.

For single molecule TIRF microscopy experiments involving mEos-SHIP1(PH-PP-C2), PLB-985 cells expressing low concentrations were identified based on 500-550 nm fluorescence emission following excitation with 488 nm light. Next, the red channel was imaged to confirm the lack of 575-625 nm emission before photoconversion. To photoconvert mEos from the green to red state, cells were globally excited with 1-3 mW of 405 light for 50-100 ms. To track association and dissociation of mEos-SHIP1(PH-PP-C2) at the plasma membrane, a 256 x 256-pixel region of interest (RIO) was recorded on an iXion Life 897 EMCCD camera at 45 frames per sec (i.e. 22 ms interval).

### Analysis of single molecule TIRF microscopy data

Single particle detection and tracking was performed using the TrackMate plugin (57) on ImageJ/Fiji as previously described (33, 48). Output files are analyzed using Prism 9 graphing software. To calculate the single molecule dwell times for fluorescently labeled SHIP1 we generated a cumulative distribution frequency (CDF) plot using the frame interval as the bin size (e.g. 22 ms). The log_10_(1-CDF) was plotted against the dwell time and fit with either a single or double exponential decay curve.

Single exponential model:

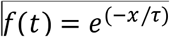

Two exponential model:

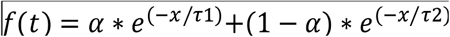

Fitting procedure initiated with a single exponential. In cases of a low-quality single exponential fit, a maximum of two populations were used. For double exponential fit, alpha (α) represents the percentage of short dwelling molecules characterized by the time constant (τ_1_). Errors reported for the single molecule dwell times represent the standard deviation from multiple technical replicates as indicated in **Table 1** and the figure legends.

To calculate the diffusion coefficients (µm^2^/sec) for membrane bound mNG-SHIP1(PH-PP-C2) in **Figure S7** and **Table 1**, we plotted probability density (i.e. frequency divided by bin size of 0.01 µm) versus step size (µm). The single molecule displacement was measured over a 12 ms time interval with ∼20,000 displacements detected per technical replicate. Step size distributions were fit to the following two species model:

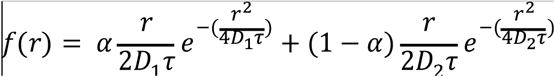

To calculate the cumulative membrane binding frequency in **Figure 5** and **Figure S6**, molecules were analyzed using the ImageJ/Fiji plugin, TrackMate. This allowed us to determine the exact frame when a mNG-SHIP1 molecule was first detected, which was then scored as a single membrane binding event. Although the program continues to track every molecule until membrane dissociation or photobleaching occurs, only new mNG-SHIP1 membrane docking events are scored and used to calculate the cumulative membrane binding frequency. All binding events for a single video were summed over a 30 second period, which represents cumulative membrane binding for a single mNG-SHIP1 concentration between 200-800 pM (**Figure 5D** and **Figure S6**).

To quantify the fraction of membrane localized mNG-tagged SHIP1 monomers and dimers (**Figure 6**), we wrote custom Python code extract the particle brightness in the second frame of each single molecule trajectory. To ensure emitted photons were collects for the full duration of a 52 ms exposure, we measured the particle brightness in the second frame of tracks that were ≥ 156 ms (i.e. ≥ 3 frames). This approach is necessary because a molecule can associate or dissociate in the middle of the first or last frame of a trajectory. This results in fewer emitted photons and a lower observed molecular brightness. Particles were identified using TrackMate (ImageJ) using an object diameter of 7 pixels. The particle brightnesses reported in **Figure 6H** represent the maximum pixel intensity for each mNG-SHIP1 molecule detected.

## REFERENCES

1. Di Paolo, G., and De Camilli, P. (2006) Phosphoinositides in cell regulation and membrane dynamics. Nature. 443, 651–657

2. Hammond, G. R. V., and Burke, J. E. (2020) Novel roles of phosphoinositides in signaling, lipid transport, and disease. Curr Opin Cell Biol. 63, 57–67

3. Balla, T. (2013) Phosphoinositides: tiny lipids with giant impact on cell regulation. Physiol. Rev. 93, 1019–1137

4. Servant, G., Weiner, O. D., Herzmark, P., Balla, T., Sedat, J. W., and Bourne, H. R. (2000) Polarization of Chemoattractant Receptor Signaling During Neutrophil Chemotaxis. Science. 287, 1037–1040

5. Stephens, L. R., Hughes, K. T., and Irvine, R. F. (1991) Pathway of phosphatidylinositol(3,4,5)-trisphosphate synthesis in activated neutrophils. Nature. 351, 33–39

6. Vadas, O., Burke, J. E., Zhang, X., Berndt, A., and Williams, R. L. (2011) Structural Basis for Activation and Inhibition of Class I Phosphoinositide 3-Kinases. Sci. Signal. 4, re2–re2

7. Wang, F., Herzmark, P., Weiner, O. D., Srinivasan, S., Servant, G., and Bourne, H. R. (2002) Lipid products of PI(3)Ks maintain persistent cell polarity and directed motility in neutrophils. Nature Cell Biology. 4, 513–518

8. Damen, J. E., Liu, L., Rosten, P., Humphries, R. K., Jefferson, A. B., Majerus, P. W., and Krystal, G. (1996) The 145-kDa protein induced to associate with Shc by multiple cytokines is an inositol tetraphosphate and phosphatidylinositol 3,4,5-triphosphate 5-phosphatase. PNAS. 93, 1689–1693

9. Goulden, B. D., Pacheco, J., Dull, A., Zewe, J. P., Deiters, A., and Hammond, G. R. V. (2019) A high-avidity biosensor reveals plasma membrane PI(3,4)P2 is predominantly a class I PI3K signaling product. Journal of Cell Biology. 218, 1066–1079

10. Pesesse, X., Deleu, S., De Smedt, F., Drayer, L., and Erneux, C. (1997) Identification of a Second SH2-Domain-Containing Protein Closely Related to the Phosphatidylinositol Polyphosphate 5-Phosphatase SHIP. Biochemical and Biophysical Research Communications. 239, 697–700

11. Belliveau, N. M., Footer, M. J., Akdoğan, E., Van Loon, A. P., Collins, S. R., and Theriot, J. A. (2023) Whole-genome screens reveal regulators of differentiation state and context-dependent migration in human neutrophils. Nat Commun. 14, 5770

12. Collins, S. R., Yang, H. W., Bonger, K. M., Guignet, E. G., Wandless, T. J., and Meyer, T. (2015) Using light to shape chemical gradients for parallel and automated analysis of chemotaxis. Molecular Systems Biology. 11, 804

13. Mondal, S., Subramanian, K. K., Sakai, J., Bajrami, B., and Luo, H. R. (2012) Phosphoinositide lipid phosphatase SHIP1 and PTEN coordinate to regulate cell migration and adhesion. MBoC. 23, 1219–1230

14. Nishio, M., Watanabe, K., Sasaki, J., Taya, C., Takasuga, S., Iizuka, R., Balla, T., Yamazaki, M., Watanabe, H., Itoh, R., Kuroda, S., Horie, Y., Förster, I., Mak, T. W., Yonekawa, H., Penninger, J. M., Kanaho, Y., Suzuki, A., and Sasaki, T. (2007) Control of cell polarity and motility by the PtdIns(3,4,5)P3 phosphatase SHIP1. Nat Cell Biol. 9, 36–44

15. Helgason, C. D., Damen, J. E., Rosten, P., Grewal, R., Sorensen, P., Chappel, S. M., Borowski, A., Jirik, F., Krystal, G., and Humphries, R. K. (1998) Targeted disruption of SHIP leads to hemopoietic perturbations, lung pathology, and a shortened life span. Genes Dev. 12, 1610–1620

16. Chou, V., Pearse, R. V., Aylward, A. J., Ashour, N., Taga, M., Terzioglu, G., Fujita, M., Fancher, S. B., Sigalov, A., Benoit, C. R., Lee, H., Lam, M., Seyfried, N. T., Bennett, D. A., De Jager, P. L., Menon, V., and Young-Pearse, T. L. (2023) INPP5D regulates inflammasome activation in human microglia. Nat Commun. 14, 7552

17. Hansen, D. V., Hanson, J. E., and Sheng, M. (2018) Microglia in Alzheimer’s disease. Journal of Cell Biology. 217, 459–472

18. Terzioglu, G., and Young-Pearse, T. L. (2023) Microglial function, INPP5D/SHIP1 signaling, and NLRP3 inflammasome activation: implications for Alzheimer’s disease. Mol Neurodegeneration. 18, 89

19. Kavanaugh, W. M., Pot, D. A., Chin, S. M., Deuter-Reinhard, M., Jefferson, A. B., Norris, F. A., Masiarz, F. R., Cousens, L. S., Majerus, P. W., and Williams, L. T. (1996) Multiple forms of an inositol polyphosphate 5-phosphatase form signaling complexes with Shc and Grb2. Current Biology. 6, 438–445

20. Zhou, H., Yue, X., Wang, Z., Li, S., Zhu, J., Yang, Y., and Liu, M. (2021) Expression, purification and characterization of the RhoA-binding domain of human SHIP2 in E.coli. Protein Expr Purif. 180, 105821

21. Bradshaw, W. J., Kennedy, E. C., Moreira, T., Smith, L. A., Chalk, R., Katis, V. L., Benesch, J. L. P., Brennan, P. E., Murphy, E. J., and Gileadi, O. (2024) Regulation of inositol 5-phosphatase activity by the C2 domain of SHIP1 and SHIP2. Structure. 32, 453-466.e6

22. Le Coq, J., López Navajas, P., Rodrigo Martin, B., Alfonso, C., and Lietha, D. (2021) A new layer of phosphoinositide-mediated allosteric regulation uncovered for SHIP2. FASEB j. 10.1096/fj.202100561R

23. Le Coq, J., Camacho-Artacho, M., Velázquez, J. V., Santiveri, C. M., Gallego, L. H., Campos-Olivas, R., Dölker, N., and Lietha, D. (2017) Structural basis for interdomain communication in SHIP2 providing high phosphatase activity. eLife. 6, e26640

24. Waddell, G. L., Drew, E. E., Rupp, H. P., and Hansen, S. D. (2023) Mechanisms controlling membrane recruitment and activation of the autoinhibited SHIP1 inositol 5-phosphatase. Journal of Biological Chemistry. 10.1016/j.jbc.2023.105022

25. Chung, J. K., Nocka, L. M., Decker, A., Wang, Q., Kadlecek, T. A., Weiss, A., Kuriyan, J., and Groves, J. T. (2019) Switch-like activation of Bruton’s tyrosine kinase by membrane-mediated dimerization. PNAS. 116, 10798–10803

26. Duewell, B. R., Wilson, Naomi E., N. E., Bailey, G. E., Peabody, S. E., and Hansen, S. D. (2023) Molecular dissection of PI3Kβ synergistic activation by receptor tyrosine kinases, GβGγ, and Rho-family GTPases. eLife. 10.7554/eLife.88991

27. Masson, G. R., Jenkins, M. L., and Burke, J. E. (2017) An overview of hydrogen deuterium exchange mass spectrometry (HDX-MS) in drug discovery. Expert Opin Drug Discov. 12, 981–994

28. Jumper, J., Evans, R., Pritzel, A., Green, T., Figurnov, M., Ronneberger, O., Tunyasuvunakool, K., Bates, R., Žídek, A., Potapenko, A., Bridgland, A., Meyer, C., Kohl, S. A. A., Ballard, A. J., Cowie, A., Romera-Paredes, B., Nikolov, S., Jain, R., Adler, J., Back, T., Petersen, S., Reiman, D., Clancy, E., Zielinski, M., Steinegger, M., Pacholska, M., Berghammer, T., Bodenstein, S., Silver, D., Vinyals, O., Senior, A. W., Kavukcuoglu, K., Kohli, P., and Hassabis, D. (2021) Highly accurate protein structure prediction with AlphaFold. Nature. 596, 583–589

29. Varadi, M., Anyango, S., Deshpande, M., Nair, S., Natassia, C., Yordanova, G., Yuan, D., Stroe, O., Wood, G., Laydon, A., Žídek, A., Green, T., Tunyasuvunakool, K., Petersen, S., Jumper, J., Clancy, E., Green, R., Vora, A., Lutfi, M., Figurnov, M., Cowie, A., Hobbs, N., Kohli, P., Kleywegt, G., Birney, E., Hassabis, D., and Velankar, S. (2022) AlphaFold Protein Structure Database: massively expanding the structural coverage of protein-sequence space with high-accuracy models. Nucleic Acids Research. 50, D439–D444

30. Abramson, J., Adler, J., Dunger, J., Evans, R., Green, T., Pritzel, A., Ronneberger, O., Willmore, L., Ballard, A. J., Bambrick, J., Bodenstein, S. W., Evans, D. A., Hung, C.-C., O’Neill, M., Reiman, D., Tunyasuvunakool, K., Wu, Z., Žemgulytė, A., Arvaniti, E., Beattie, C., Bertolli, O., Bridgland, A., Cherepanov, A., Congreve, M., Cowen-Rivers, A. I., Cowie, A., Figurnov, M., Fuchs, F. B., Gladman, H., Jain, R., Khan, Y. A., Low, C. M. R., Perlin, K., Potapenko, A., Savy, P., Singh, S., Stecula, A., Thillaisundaram, A., Tong, C., Yakneen, S., Zhong, E. D., Zielinski, M., Žídek, A., Bapst, V., Kohli, P., Jaderberg, M., Hassabis, D., and Jumper, J. M. (2024) Addendum: Accurate structure prediction of biomolecular interactions with AlphaFold 3. Nature. 636, E4

31. Nalefski, E. A., and Falke, J. J. (1996) The C2 domain calcium-binding motif: Structural and functional diversity. Protein Science. 5, 2375–2390

32. Lorent, J. H., Levental, K. R., Ganesan, L., Rivera-Longsworth, G., Sezgin, E., Doktorova, M., Lyman, E., and Levental, I. (2020) Plasma membranes are asymmetric in lipid unsaturation, packing and protein shape. Nat Chem Biol. 16, 644–652

33. Hansen, S. D., Lee, A. A., Duewell, B. R., and Groves, J. T. (2022) Membrane-mediated dimerization potentiates PIP5K lipid kinase activity. eLife. 11, e73747

34. Chung, J. K., Lee, Y. K., Denson, J.-P., Gillette, W. K., Alvarez, S., Stephen, A. G., and Groves, J.T. (2018) K-Ras4B Remains Monomeric on Membranes over a Wide Range of Surface Densities and Lipid Compositions. Biophysical Journal. 114, 137–145

35. Knight, J. D., Lerner, M. G., Marcano-Velázquez, J. G., Pastor, R. W., and Falke, J. J. (2010) Single Molecule Diffusion of Membrane-Bound Proteins: Window into Lipid Contacts and Bilayer Dynamics. Biophysical Journal. 99, 2879–2887

36. Kim, A. S., Kakalis, L. T., Abdul-Manan, N., Liu, G. A., and Rosen, M. K. (2000) Autoinhibition and activation mechanisms of the Wiskott-Aldrich syndrome protein. Nature. 404, 151–158

37. Prehoda, K. E. (2000) Integration of Multiple Signals Through Cooperative Regulation of the N-WASP-Arp2/3 Complex. Science. 290, 801–806

38. Corbalan-Garcia, S., and Gómez-Fernández, J. C. (2014) Signaling through C2 domains: More than one lipid target. Biochimica et Biophysica Acta (BBA) - Biomembranes. 1838, 1536–1547

39. Jiménez, J. L., Smith, G. R., Contreras-Moreira, B., Sgouros, J. G., Meunier, F. A., Bates, P. A., and Schiavo, G. (2003) Functional Recycling of C2 Domains Throughout Evolution: A Comparative Study of Synaptotagmin, Protein Kinase C and Phospholipase C by Sequence, Structural and Modelling Approaches. Journal of Molecular Biology. 333, 621–639

40. Cho, W., and Stahelin, R. (2006) Membrane binding and subcellular targeting of C2 domains. Biochimica et Biophysica Acta (BBA) - Molecular and Cell Biology of Lipids. 1761, 838–849

41. Hejna, J. A., Saito, H., Merkens, L. S., Tittle, T. V., Jakobs, P. M., Whitney, M. A., Grompe, M., Friedberg, A. S., and Moses, R. E. (1995) Cloning and Characterization of a Human cDNA (INPPL1) Sharing Homology with Inositol Polyphosphate Phosphatases. Genomics. 29, 285–287

42. Gibson, D. G., Young, L., Chuang, R.-Y., Venter, J. C., Hutchison, C. A., and Smith, H. O. (2009) Enzymatic assembly of DNA molecules up to several hundred kilobases. Nat Methods. 6, 343–345

43. Gee, P., Lung, M. S. Y., Okuzaki, Y., Sasakawa, N., Iguchi, T., Makita, Y., Hozumi, H., Miura, Y., Yang, L. F., Iwasaki, M., Wang, X. H., Waller, M. A., Shirai, N., Abe, Y. O., Fujita, Y., Watanabe, K., Kagita, A., Iwabuchi, K. A., Yasuda, M., Xu, H., Noda, T., Komano, J., Sakurai, H., Inukai, N., and Hotta, A. (2020) Extracellular nanovesicles for packaging of CRISPR-Cas9 protein and sgRNA to induce therapeutic exon skipping. Nat Commun. 11, 1334

44. Lin, W.-C., Yu, C.-H., Triffo, S., and Groves, J. T. (2010) Supported membrane formation, characterization, functionalization, and patterning for application in biological science and technology. Curr Protoc Chem Biol. 2, 235–269

45. Mukherjee, O., Weingarten, L., Padberg, I., Pracht, C., Sinha, R., Hochdörfer, T., Kuppig, S., Backofen, R., Reth, M., and Huber, M. (2012) The SH2-domain of SHIP1 interacts with the SHIP1 C-terminus: Impact on SHIP1/Ig-α interaction. Biochimica et Biophysica Acta (BBA) - Molecular Cell Research. 1823, 206–214

46. Hansen, S. D., Huang, W. Y. C., Lee, Y. K., Bieling, P., Christensen, S. M., and Groves, J. T. (2019) Stochastic geometry sensing and polarization in a lipid kinase–phosphatase competitive reaction. Proc. Natl. Acad. Sci. U.S.A. 116, 15013–15022

47. Vacklin, H. P., Tiberg, F., and Thomas, R. K. (2005) Formation of supported phospholipid bilayers via co-adsorption with beta-D-dodecyl maltoside. Biochim. Biophys. Acta. 1668, 17–24

48. Hansen, S. D., Huang, W. Y. C., Lee, Y. K., Bieling, P., Christensen, S. M., and Groves, J. T. (2019) Stochastic geometry sensing and polarization in a lipid kinase–phosphatase competitive reaction. Proc Natl Acad Sci USA. 116, 15013–15022

49. Cordes, T., Vogelsang, J., and Tinnefeld, P. (2009) On the Mechanism of Trolox as Antiblinking and Antibleaching Reagent. J. Am. Chem. Soc. 131, 5018–5019

50. Stariha, J. T. B., Hoffmann, R. M., Hamelin, D. J., and Burke, J. E. (2021) Probing Protein-Membrane Interactions and Dynamics Using Hydrogen-Deuterium Exchange Mass Spectrometry (HDX-MS). Methods Mol Biol. 2263, 465–485

51. da Veiga Leprevost, F., Haynes, S. E., Avtonomov, D. M., Chang, H.-Y., Shanmugam, A. K., Mellacheruvu, D., Kong, A. T., and Nesvizhskii, A. I. (2020) Philosopher: a versatile toolkit for shotgun proteomics data analysis. Nat Methods. 17, 869–870

52. Dobbs, J. M., Jenkins, M. L., and Burke, J. E. (2020) Escherichia coli and Sf9 Contaminant Databases to Increase Efficiency of Tandem Mass Spectrometry Peptide Identification in Structural Mass Spectrometry Experiments. J Am Soc Mass Spectrom. 31, 2202–2209

53. Kong, A. T., Leprevost, F. V., Avtonomov, D. M., Mellacheruvu, D., and Nesvizhskii, A. I. (2017) MSFragger: ultrafast and comprehensive peptide identification in mass spectrometry-based proteomics. Nat Methods. 14, 513–520

54. Masson, G. R., Burke, J. E., Ahn, N. G., Anand, G. S., Borchers, C., Brier, S., Bou-Assaf, G. M., Engen, J. R., Englander, S. W., Faber, J., Garlish, R., Griffin, P. R., Gross, M. L., Guttman, M., Hamuro, Y., Heck, A. J. R., Houde, D., Iacob, R. E., Jørgensen, T. J. D., Kaltashov, I. A., Klinman, J. P., Konermann, L., Man, P., Mayne, L., Pascal, B. D., Reichmann, D., Skehel, M., Snijder, J., Strutzenberg, T. S., Underbakke, E. S., Wagner, C., Wales, T. E., Walters, B. T., Weis, D. D., Wilson, D. J., Wintrode, P. L., Zhang, Z., Zheng, J., Schriemer, D. C., and Rand, K. D. (2019) Recommendations for performing, interpreting and reporting hydrogen deuterium exchange mass spectrometry (HDX-MS) experiments. Nat Methods. 16, 595–602

55. Perez-Riverol, Y., Bai, J., Bandla, C., García-Seisdedos, D., Hewapathirana, S., Kamatchinathan, S., Kundu, D. J., Prakash, A., Frericks-Zipper, A., Eisenacher, M., Walzer, M., Wang, S., Brazma, A., and Vizcaíno, J. A. (2022) The PRIDE database resources in 2022: a hub for mass spectrometry-based proteomics evidences. Nucleic Acids Res. 50, D543–D552

56. Rincón, E., Rocha-Gregg, B. L., and Collins, S. R. (2018) A map of gene expression in neutrophil-like cell lines. BMC Genomics. 19, 573

57. Jaqaman, K., Loerke, D., Mettlen, M., Kuwata, H., Grinstein, S., Schmid, S. L., and Danuser, G. (2008) Robust single-particle tracking in live-cell time-lapse sequences. Nat Methods. 5, 695–702

