## Supplemental Figures for "SH2-mediated steric occlusion of the C2 domain regulates autoinhibition of SHIP1 inositol 5-phosphatase"

### Supplemental Figure 1

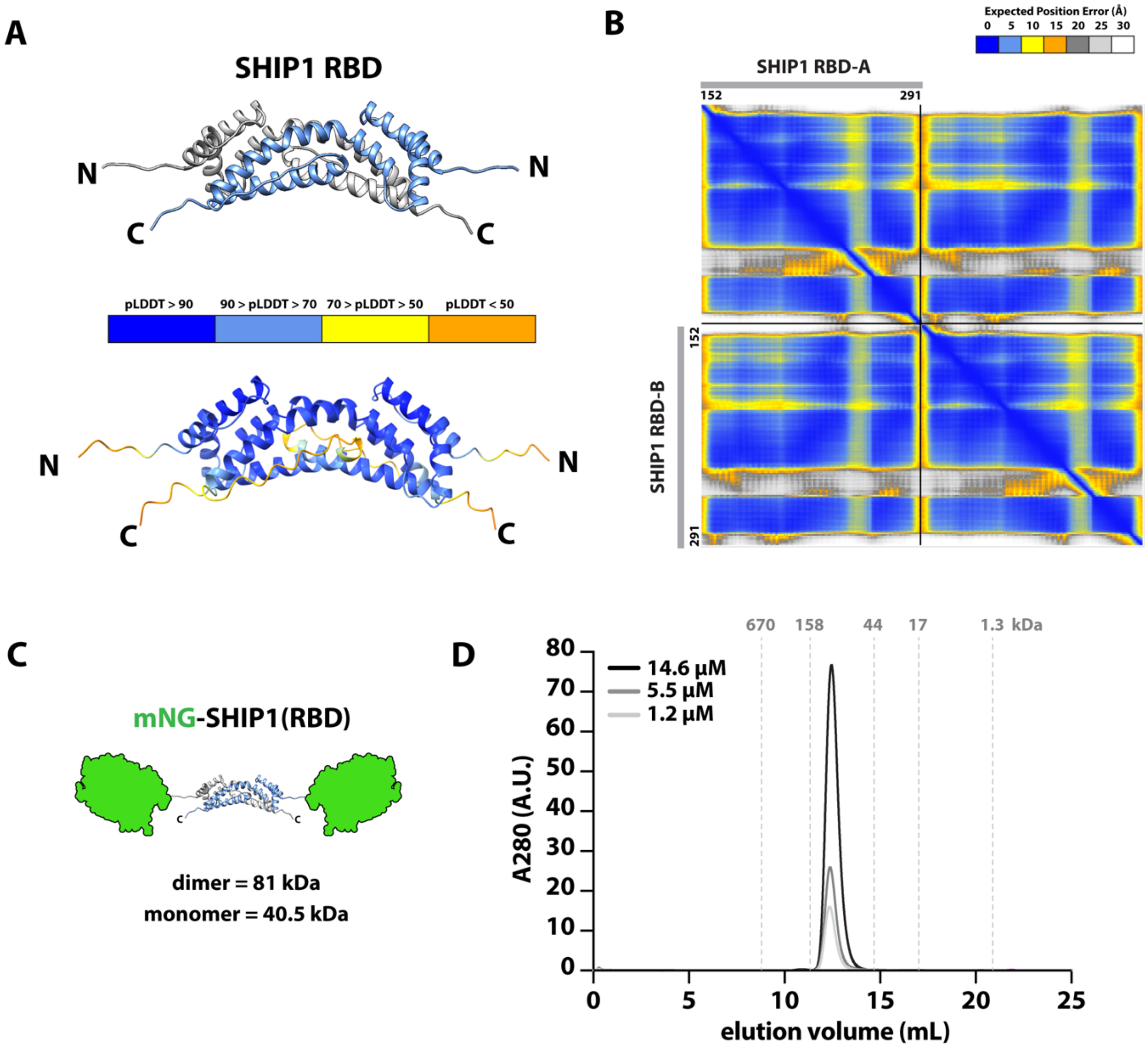


**Figure Supplement 1**

**Characterization of SHIP1 Rho binding domain (RBD)**

**(A)** AlphaFold2 Multimer prediction of the SHIP1 Rho binding domain (RBD) homodimerizing. Shown below is the model colored by predicted local distance difference (pLDDT) to show regions with high confidence (pLDDT > 90) to low confidence (pLDDT <50). **(B)** Predicted alignment error (PAE) for Alphafold2 multimer model of SHIP1 RBD dimer. Note that the PAE plot is not an inter-residue distance map or a contact map. Instead, the coloring indicates expected distance error. The color at (x, y) corresponds to the expected distance error in residue x's position (Angstroms), when the prediction are aligned on residue y (more information can be found at https://alphafold.ebi.ac.uk/). **(C)** Cartoon depiction of mNG-SHIP1(RBD) with the expected molecular weight of a dimer (81 kDa) or monomer (40.5 kDa). **(D)** Size exclusion chromatography elution profiles for varying concentration of mNG-SHIP1(RBD) injected on a Superdex200 Increase 10/300GL column (Cytiva, Cat# 28990944). Load concentration indicated in the legend. Elution profile of molecular weight standards (Bio-Rad, Cat#151-1901) indicated by grey dashed lines and kDa MW's above.

### Supplemental Figure 2

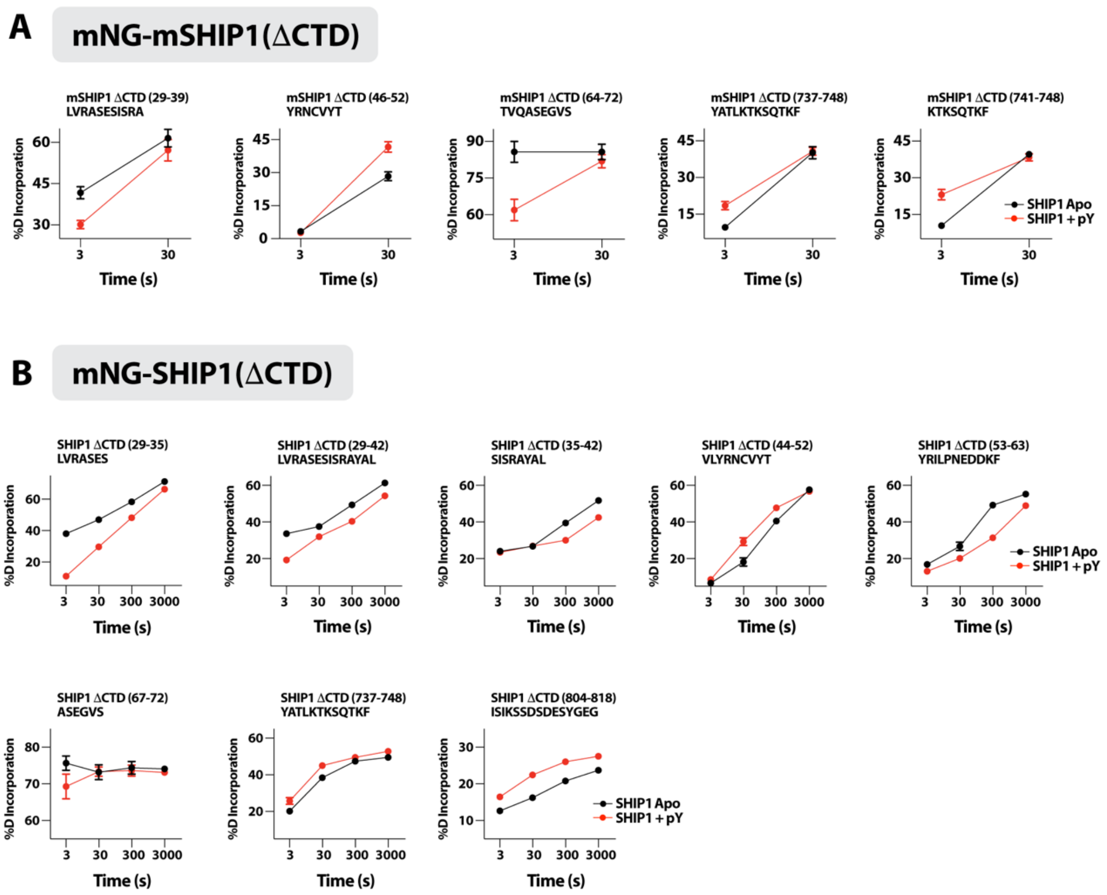


**Figure Supplement 2**
**Percent deuterium incorporation graphs for SHIP1**
Mean of the % deuterium uptake for **(A)** mNG-mSHIP1(ΔCTD) and **(B)** mNG-SHIP1(ΔCTD) peptides that showed a significant change in HDX (>0.4 Da and 5% difference, with a two-tailed t-test $p < 0.01$) across the entire deuterium exchange time course (error bars represent standard deviation, n = 3 for all timepoints).

### Supplemental Figure 3

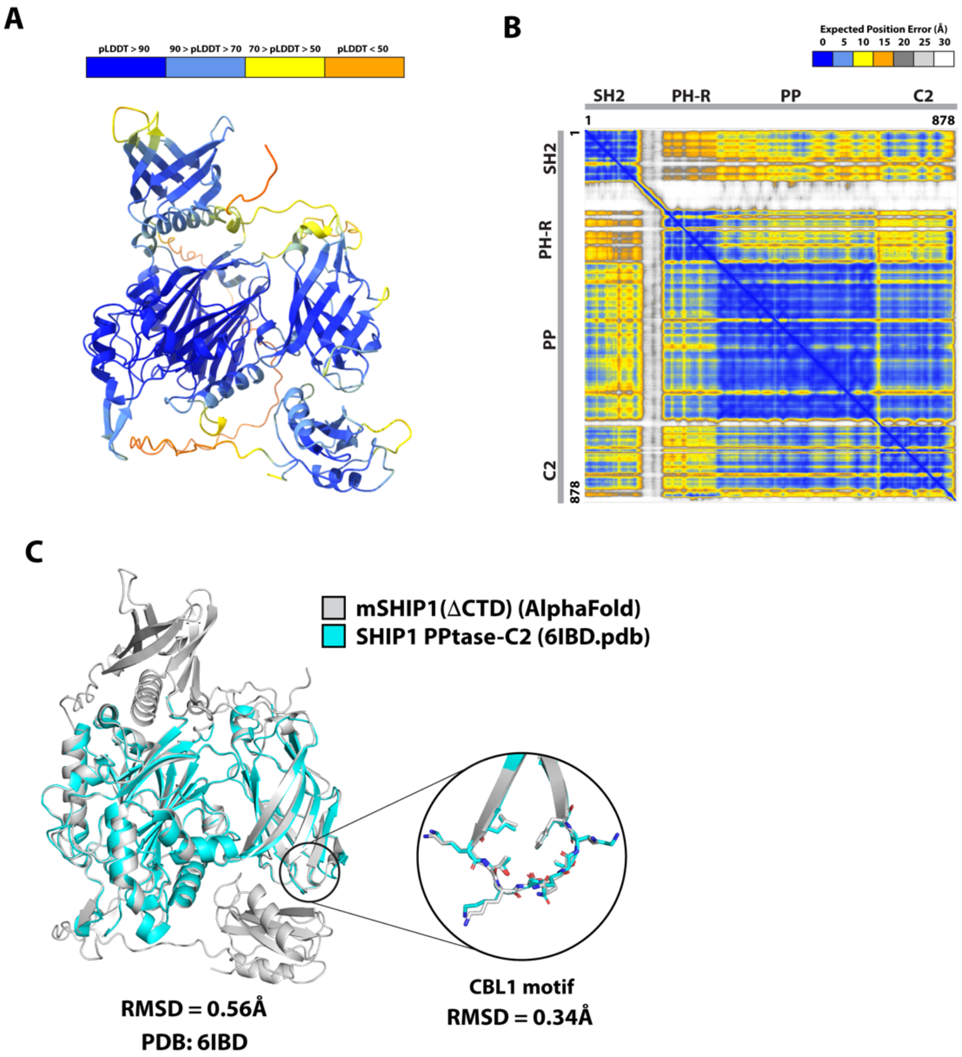


**Figure Supplement 3**

**Generation and validation of an AlphaFold model for monomeric SHIP1(ΔCTD)**

**(A)** AlphaFold2 Multimer prediction of mSHIP1(ΔCTD). Shown below is the model colored by predicted local distance difference (pLDDT) to show regions with high confidence (pLDDT > 90) to low confidence (pLDDT <50). **(B)** Predicted alignment error (PAE) for Alphafold2 multimer model of mini-SHIP1. Note that the PAE plot is not an inter-residue distance map or a contact map. Instead, the coloring indicates expected distance error. **(C)** Overlay of AlphaFold predicted structure of mSHIP1(ΔCTD) (grey) and the X-ray crystal structure human SHIP1 PPtase-C2 (397-857aa, teal). Root mean square deviation (RMSD) for the entire protein and CBL1 motif indicated under the structures.

### Supplemental Figure 4

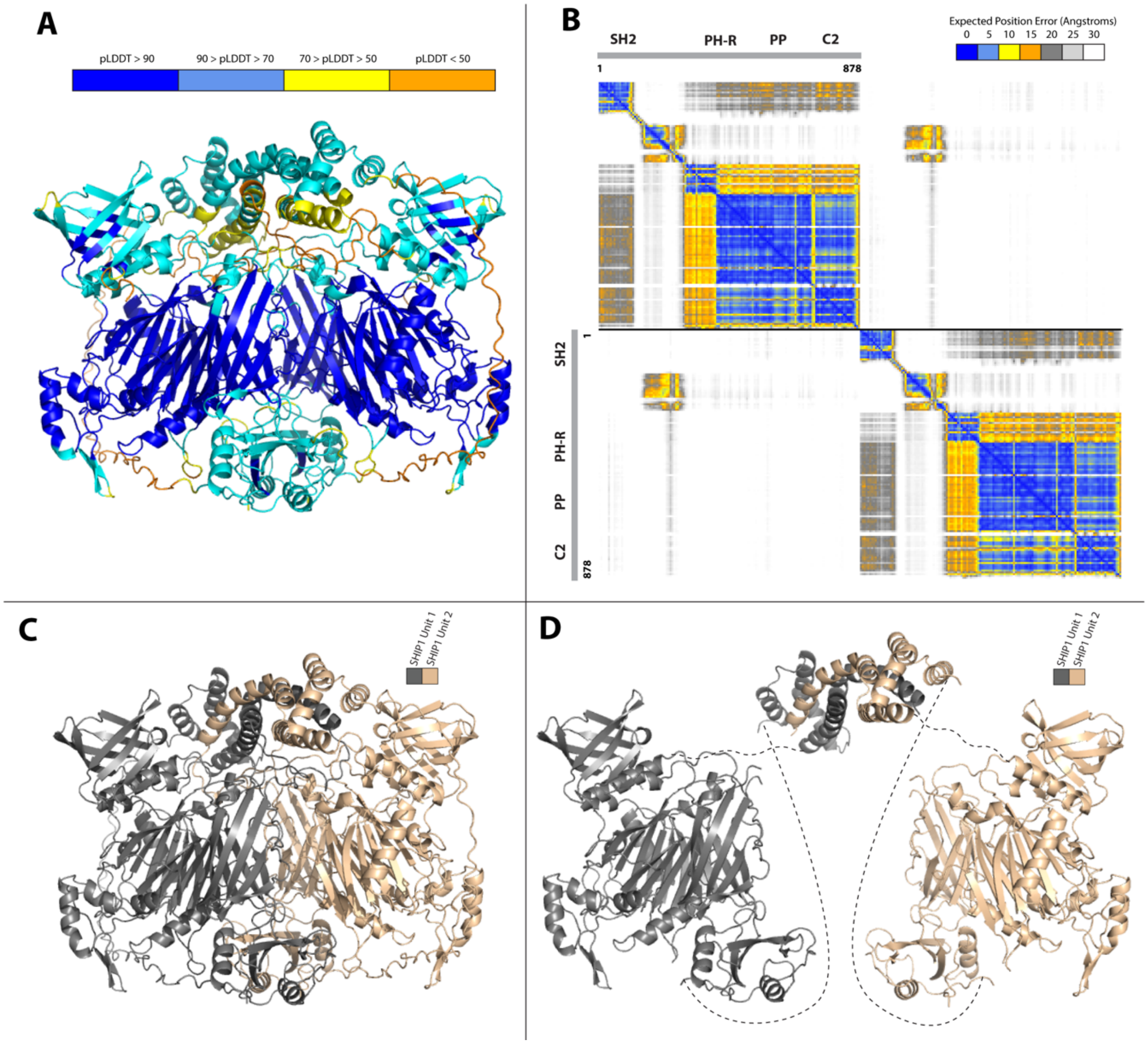


**Figure Supplement 4**

**Generation and validation of an AlphaFold model of the SHIP1(ΔCTD) dimer**

**(A)** AlphaFold 3 search of two copies of SHIP1(ΔCTD). Model is colored by pLDDT to indicate regions of high and low confidence. **(B)** Predicted alignment error (PAE) for AlphaFold3 search of two copies of SHIP1(ΔCTD). The colors indicate the predicted aligned error and are colored according to the legend. **(C)** Structure of SHIP1 dimer colored by chain to differentiate between subunits. Residues with low local confidence (pLDDT < 50) have been removed. **(D)** Modelled interfaces with a high expected position error were manually separated from each other. Low confidence interdomain contacts with the dimerization domain which had been removed from the AlphaFold model were manually annotated using dashed lines.

### Supplemental Figure 5

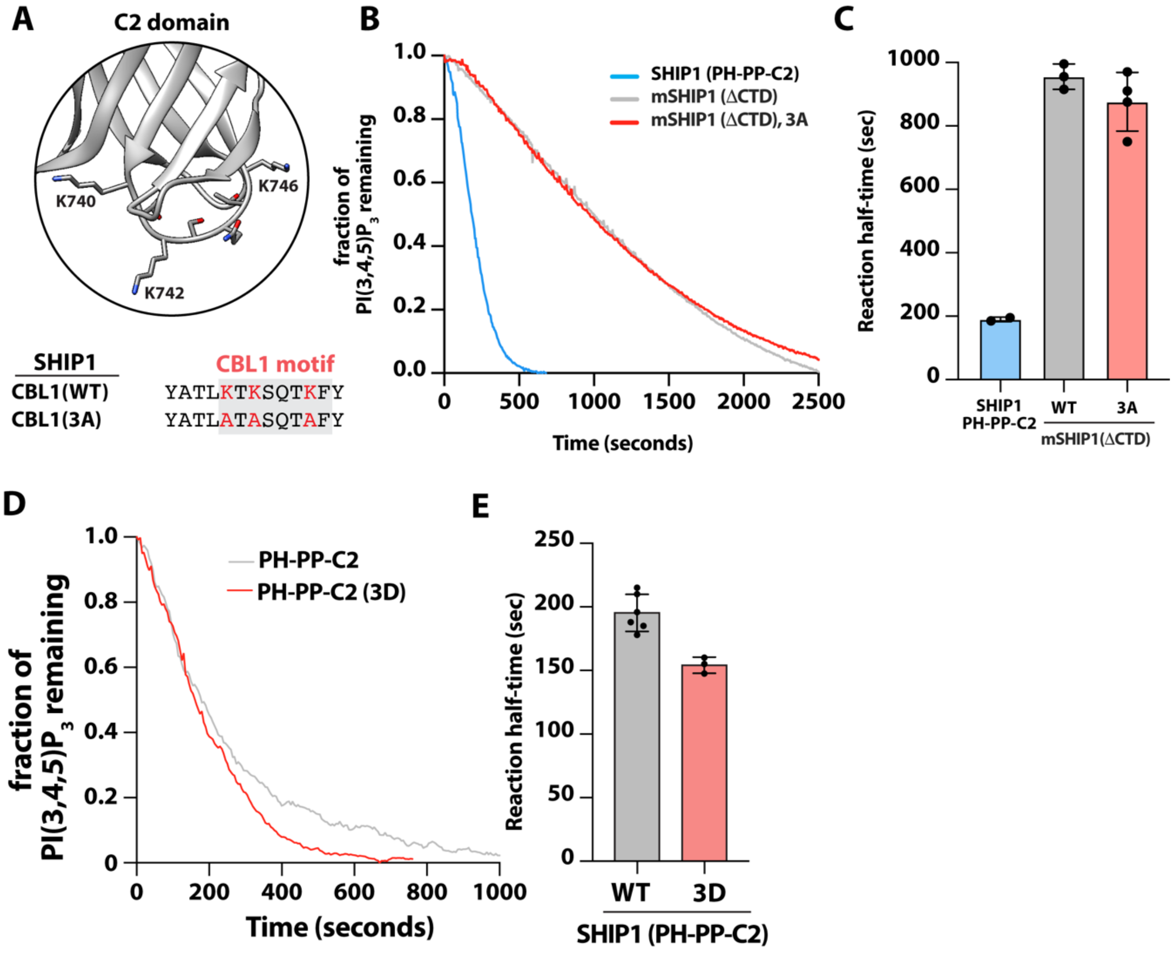


**Figure Supplement 5**

**SHIP1 CBL1 motif mutant lipid phosphatase activity measurements**

**(A)** Structural model of human SHIP1 CBL1 motif with indicated lysine residues (6IBD.pdb). Sequence alignment comparing human SHIP1 CBL1 motif, wild type and mutant (K741A/K743A/K747A, denoted 3A) sequence. **(B)** Charge neutralization (3A) in the CBL1 motif of mini-SHIP1 does not enhance lipid phosphatase activity. Kinetic traces of phosphatase activity measured in the presence of 20 nM mNG-mSHIP1(ΔCTD, WT or 3A) and 20 nM mNG-SHIP1 (PH-PP-C2). **(C)** Quantification of reaction half-times measured in the presence of 20 nM mNG-SHIP1(PH-PP-C2), 20 nM mNG-mSHIP1(ΔCTD), or 20 nM mNG-mSHIP1(ΔCTD, 3A). Bars equal to the mean reaction half-times (N= 3 technical replicates). Errors equal standard deviation. **(D)** Phosphatase measurements of 20 nM mNG-SHIP1(PH-PP-C2) or mNG-SHIP1 CBL1 motif mutants. **(E)** Quantification of reaction half-times of 20 nM mNG-SHIP1(PH-PP-C2) and 20 nM mNG- SHIP1(PH-PP-C2, K741D/K743D/K747D; denoted 3D). Bars are equal to the mean reaction half-times (N = 3-6 technical replicate per construct). Student t-test comparing SHIP1(PH-PP-C2), WT and 3D, produced a p-value = 0.003. (**B-E**) Dephosphorylation of PI(3,4,5)$P_3$ was monitored in the presence of 20 nM AF555-SNAP-Btk. Initial membrane composition: 2% PI(3,4,5)$P_3$, 98% DOPC.

### Supplemental Figure 6

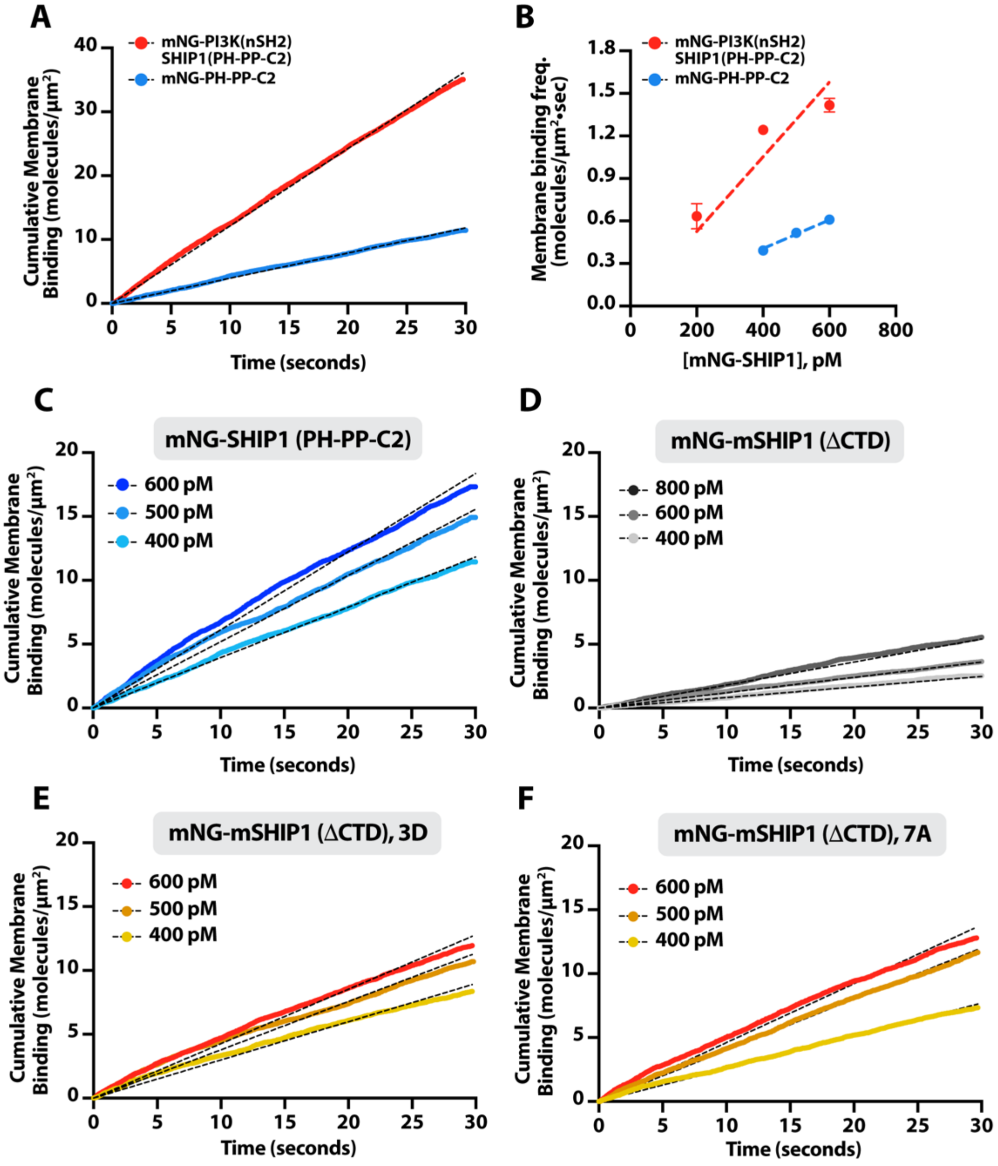


**Figure Supplement 6**
**Single molecule membrane binding frequency measurements**
**(A)** Cumulative binding events measured by TIRF-M in the presence of 400 pM mNG-SHIP1 (PH-PP-C2) and 400 pM mNG-PI3K(SH2)-SHIP1(PH-PP-C2). **(B)** Membrane binding frequency ($k_{ON}$) calculated from measuring cumulative binding events across the following protein concentrations: 400-600 pM mNG-SHIP1 (PH-PP-C2) and 200-600 pM mNG-PI3K(SH2)-SHIP1(PH-PP-C2). Linear regression yielded the following $k_{ON}$ values: 1.0 $nM^{-1}$•$\mu m^{-2}$•$sec^{-1}$ mNG-SHIP1 (PH-PP-C2) and 2.6 $nM^{-1}$•$\mu m^{-2}$•$sec^{-1}$ mNG-PI3K(SH2)-SHIP1(PH-PP-C2). Cumulative membrane binding events measured in the presence of **(C)** 400-600 pM mNG-SHIP1 (PH-PP-C2), **(D)** 400-800 pM mNG-mSHIP1(ΔCTD), **(E)** 400-600 pM mNG-mSHIP1(ΔCTD, 3D), or **(F)** 400-600 pM mNG-mSHIP1(ΔCTD, 7A). **(A-F)** Membrane composition: 2% $PI(3,4,5)P_3$, 20% DOPS, 78% DOPC.

### Supplemental Figure 7

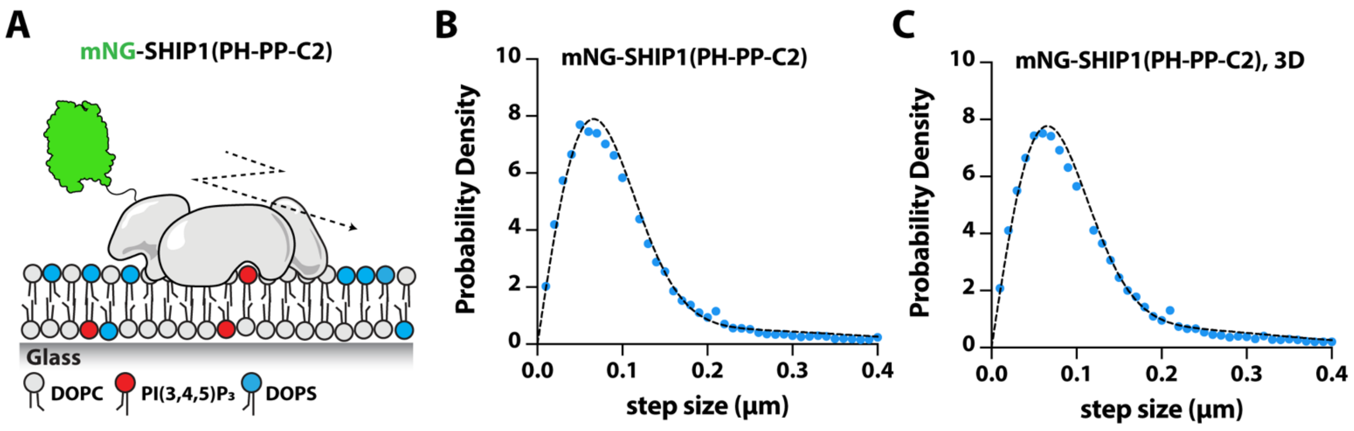


**Figure Supplement 7**

**CBL1 motif mutations do not alter membrane diffusivity of mNG-SHIP1(PH-PP-C2)**

**(A)** Cartoon schematic showing purified mNG-SHIP1(PH-PP-C2) bound to a supported lipid bilayer containing PI(3,4,5)$P_3$, DOPS, and DOPC. **(B-C)** Step size distributions (or displacement) measured in vitro in the presence of 200 pM mNG-SHIP1(PH-PP-C2) or 200 pM mNG-SHIP1(PH-PP-C2, 3D). The single molecule displacement (µm) was measured between each frame with 12 ms time intervals. **(B-C)** Membrane composition: 2% PI(3,4,5)$P_3$, 20% DOPS, 78% DOPC.

#### Table S1

| Protein Data Set | mSHIP1(ΔCTD) Apo | mSHIP1(ΔCTD) Apo + 40 µM pY peptide |
|---|---|---|
| HDX reaction details | % $D_2O$ = 66.66%<br>pH(read) = 7.5<br>Temp = 18ºC | % $D_2O$ = 66.66%<br>pH(read) = 7.5<br>Temp = 18ºC |
| HDX time course | 3s, 300s | 3s, 300s |
| HDX controls | N/A | N/A |
| Back-exchange | Corrected based off % D2O | Corrected based off % D2O |
| Number of unique peptides | 112 | 112 |
| Sequence coverage | 86.3 % | 86.3 % |
| Average peptide length/redundancy | length = 15.2<br>redundancy = 2.23 | length = 15.2<br>redundancy = 2.23 |
| Replicates | 3 | 3 |
| Repeatability | Average Stdev = 1.2% | Average Stdev = 1.0% |
| Significant difference in HDX | > 4.5 % and > 0.45 Da and unpaired t-test < 0.01 | > 4.5 % and > 0.45 Da and unpaired t-test < 0.01 |

**Table S1**
**HDX-MS data and statistics collected for mSHIP1(ΔCTD)**
The data analysis statistics for all HDX-MS experiments shown in table were performed and presented according to published guidelines (51). Results are presented as relative levels of deuterium incorporation and the only control for back exchange was the level of deuterium present in the buffer. Differences in exchange in a peptide were considered significant if they met all three of the following criteria: ≥5% change in exchange, ≥0.4 Da difference in exchange, and a p value <0.01 using a two tailed student t-test for mSHIP1(ΔCTD) experiment. The mass spectrometry proteomics data have been deposited to the ProteomeXchange Consortium via the PRIDE partner repository (52) with the dataset PXD061719 for mSHIP1(ΔCTD) data.

**Table S2**

| Protein Data Set | SHIP1 (ΔCTD) Apo | SHIP1 (ΔCTD) Apo + 40 μM pY peptide |
|---|---|---|
| HDX reaction details | % $D_2O$ = 70.7%<br>pH(read) = 7.5<br>Temp = 20ºC | % $D_2O$ = 70.7%<br>pH(read) = 7.5<br>Temp = 20ºC |
| HDX time course | 3s, 30s, 300s, 3000s | 3s, 30s, 300s, 3000s |
| HDX controls | N/A | N/A |
| Back-exchange | Corrected based off % D2O | Corrected based off % D2O |
| Number of unique peptides | 156 | 156 |
| Sequence coverage | 86 % | 86 % |
| Average peptide length/redundancy | length = 15.5<br>redundancy = 2.1 | length = 15.5<br>redundancy = 2.1 |
| Replicates | 3 | 3 |
| Repeatability | Average Stdev = 1.1% | Average Stdev = 0.9% |
| Significant difference in HDX | > 5 % and > 0.4 Da and unpaired t-test < 0.01 | > 5 % and > 0.4 Da and unpaired t-test < 0.01 |

**Table S2**
**HDX-MS data and statistics collected for SHIP1(1-878aa, ΔCTD)**
The data analysis statistics for all HDX-MS experiments shown in table were performed and presented according to published guidelines (51). Results are presented as relative levels of deuterium incorporation and the only control for back exchange was the level of deuterium present in the buffer. Differences in exchange in a peptide were considered significant if they met all three of the following criteria: ≥5% change in exchange, ≥0.4 Da difference in exchange, and a p value <0.01 using a two tailed student t-test for SHIP1(ΔCTD) experiment. The mass spectrometry proteomics data have been deposited to the ProteomeXchange Consortium via the PRIDE partner repository (52) with the dataset identifier PXD058704 for SHIP1(ΔCTD) data.
